# Reconciling fast Hepatitis B evolutionary rates with ancient co-divergence

**DOI:** 10.64898/2026.06.05.730483

**Authors:** Philippe Lemey, Xiang Ji, Bram Vrancken, Magdalini Bletsa, Pratyusa Datta, Liana E. Kafetzopoulou, Jonathon C.O. Mifsud, Guy Baele, Mahmoud Reza Pourkarim, Livia Patrono, Sebastien Calvignac-Spencer, Ludovic Orlando, Paul Bastide, Stephane Guindon, Darren Martin, Marc A. Suchard

## Abstract

Estimating evolutionary rates and divergence times for hepatitis B virus (HBV) has long been complicated by conflicting calibration approaches and extensive rate variation. To unlock the full potential of ancient and modern HBV genomic data, we develop a Bayesian mixed-effects molecular clock model that accounts for various sources of rate variation including time-dependent rate decay. Our analyses reveal a pronounced decline in evolutionary rates over time, reconciling HBV divergence estimates with human migration events across both deep and more recent timescales. We show that HBV spread into Europe through both Neolithic farming expansions and later steppe migrations, paralleling patterns proposed for Indo-European language origins. Phylogeographic reconstructions suggest that the Neolithic-associated lineage dispersed at approximately 1 km/year, consistent with archaeological estimates, while genotype D expanded during the Bronze Age at an almost threefold higher rate, plausibly driven by technological innovations underlying steppe expansions. Historical overlap between these lineages facilitated recombination, giving rise to genotype E, which has become a dominant HBV genotype in Africa. These findings demonstrate that ancient viral genomes, when analyzed with models capturing complex rate dynamics, provide a powerful lens on human prehistory and the processes shaping pathogen diversity.

## Introduction

Reconstructions of time-scaled evolutionary histories are central to our understanding of viral evolution. Timescales of outbreaks or epidemic histories spanning several months to a few centuries are often inferred from heterochronous sequence data, relying on the divergence that accumulates over the sampling time range in rapidly evolving viruses. Naïve extrapolations based on fast evolutionary rates are however problematic for deep viral evolutionary histories (Düx *et al*., 2020; Wertheim and Kosakovsky Pond, 2011). Therefore, efforts to estimate deep divergence times frequently resort to biogeographic or spatial information, or to evidence of virus-host co-evolution (Rector *et al*., 2007; Saxenhofer *et al*., 2017; Switzer *et al*., 2005). Unfortunately, these different approaches often yield conflicting estimates (Saxenhofer *et al*., 2017; Wertheim and Worobey, 2009), leaving a disconnect between short- and long-term timescales and a gap in our understanding of viral evolutionary history.

Hepatitis B virus (HBV) has become a textbook example of conflicting evolutionary rate and divergence time estimates. While classified as a DNA virus, HBV replicates via reverse transcription of a pregenomic RNA, a mechanism shared with retroviruses that can lead to relatively high evolutionary rates due to error-prone replication. However, a sample of estimates from earlier studies illustrates HBV evolutionary rate estimates differing by up to four orders of magnitude, covering a substantial range of the evolutionary rate variation across many different RNA and DNA viruses (Figure 1). Different factors may explain this variation, including rate differences within and between hosts (e.g. Vrancken *et al*. (2017)) or across clinical phenotypes (e.g. Lim *et al*. (2007)), as well as the specific genome regions analyzed. In addition, the choice of calibration appears to be particularly important, as reflected in widely varying HBV divergence time estimates Figure 1). While Bayesian dated tip molecular clock estimation resulted in time to the most recent common ancestor (tM-RCA) estimates of 2 000-4 000 years for human HBV (Zhou and Holmes, 2007), the use of internal node calibrations based on historical human migration dated the same tMRCA back to 22 000-47 100 years before present (BP) (Paraskevis *et al*., 2013). Applications of dated tip models to HBV have been criticised because of model misspecification and inadequate sampling time information (Bouckaert *et al*., 2013), but internal node calibrations based on co-divergence can also be questioned as they rely on assumptions that are difficult to ascertain.

**FIG. 1.**
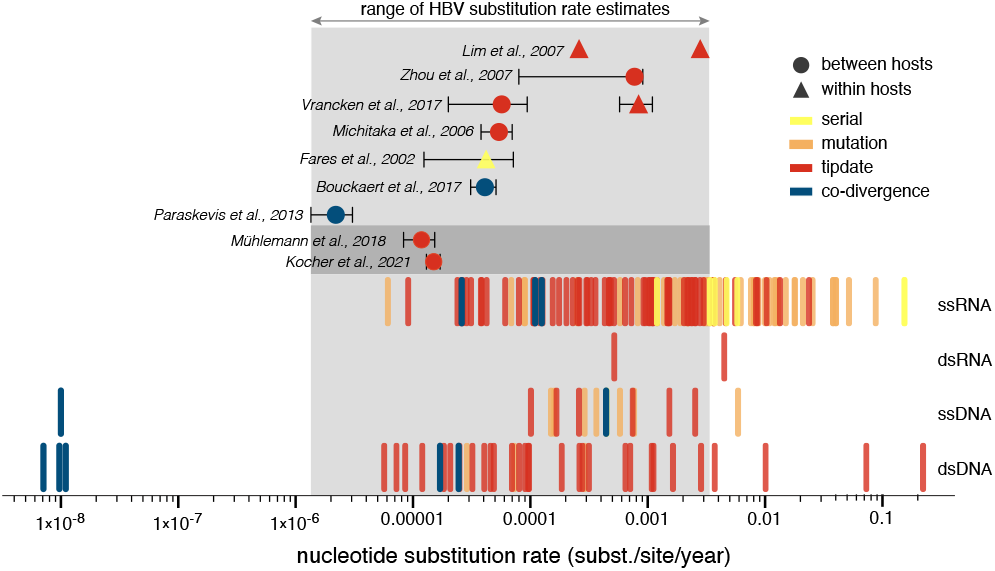
Evolutionary rate estimates for HBV and other viruses. The rug plot in the lower part of the figure summarizes mean evolutionary rate estimates collected by Duchêne *et al*. (2014b) for different viral samples and methods, including rates obtained through mutation experiments (‘mutation’), rates estimated by sampling a host or group of hosts over a timeframe of a few months to five years (‘serial’), phylogenetic estimates using a dated-tip model (‘tip-date’) and rates estimated using internal node calibration based on co-divergence assumptions (‘co-divergence’). To demonstrate the variability in HBV evolutionary rates in the upper part of the figure, we depict a sample of estimates (mean and uncertainty if available) from the literature, including estimates based on co-divergence calibrations (Bouckaert *et al*., 2017; Paraskevis *et al*., 2013), serial sampling within hosts (Fares and Holmes, 2002), and phylogenetic dated tip estimation within hosts (Lim *et al*., 2007), between hosts (Michitaka *et al*., 2006; Zhou and Holmes, 2007), or combining both (Vrancken *et al*., 2017). The range of these estimates is highlighted with a grey box. We also include dated tip estimates for analyses including ancient genomes (Kocher *et al*. (2021); Mühlemann *et al*. (2018a), highlighted with a darker grey box.

The longstanding HBV dating controversy has been further complicated by recent ancient DNA studies (Barquera *et al*., 2020; Kocher *et al*., 2021; Krause-Kyora *et al*., 2018; Mühlemann *et al*., 2018a; Patterson Ross *et al*., 2018). In 2018, Mühlemann *et al*. (2018a) discovered 12 full or partial ancient HBV genomes in shotgun sequencing libraries from human samples that are between *∼* 800 and *∼* 4 500 years old. More recently, Kocher *et al*. (2021) greatly expanded bthis effort by generating ancient HBV genomes from 137 individuals from Eurasia and the Americas dating back from *∼* 400 and *∼* 10 500 years BP. The inclusion of these ancient genomes in dated tip inference resulted in slower evolutionary rate estimates compared to dated tip estimates using modern genomes, but not as slow as the estimates obtained using ancient human migration calibrations (Figure 1, Paraskevis *et al*. (2013)). Importantly, divergence times estimated from the ancient genomes were also not compatible with previously used calibrations based on human migration (Mühlemann *et al*., 2018a), invalidating proposed theories about the origin and spread of HBV.

The ancient HBV genome data also revealed that differences in evolutionary rate estimates are not necessarily due to dated tip versus internal node calibrations, but due to the timescale of these calibrations. Specifically, dated tips encompassing deep timescales result in slow evolutionary rates (Kocher *et al*., 2021; Mühlemann *et al*., 2018a) while internal node calibrations based on relatively recent human migrations result in comparatively faster evolutionary rates (Bouckaert *et al*., 2017) (Figure 1). The fact that evolutionary rates depend on the timescale of measurement is in line with the time-dependent rate (TDR) phenomenon that is now well-established across different viruses (Aiewsakun and Katzourakis, 2016; Duchêne *et al*., 2014a). Kocher *et al*. (2021) explored a TDR molecular clock model (Membrebe *et al*., 2019) in their HBV evolutionary analyses, but found the uncorrelated relaxed clock model (Drummond *et al*., 2006) to provide a better model fit despite the fact that the TDR model recovered more consistent deep branching patterns. This suggests that extensive rate variation among branches may overwhelm the TDR signal, and that molecular clock models are needed to accommodate various sources of rate variation. Several aspects further complicate accurate HBV divergence estimation including a compact genome organization with overlapping reading frames, recombination and uncertain time stamps for ancient genomes. Here, we confront these challenges using novel Bayesian evolutionary inference methodology and demonstrate that this can reconcile long-standing conflicts about HBV evolutionary history and reveal further associations with historical human migrations.

## Materials and Methods

### Data set

We compiled a complete genome data set that includes all the ancient HBV genomes analyzed by Kocher *et al*. (2021) (*n* = 124) dated between *∼* 300 and *∼* 10 600 years BP. This comprises their newly obtained ancient genomes (*n* = 105, not including the low coverage genomes, Kocher *et al*. (2021)) and ancient genomes previously generated by Mühlemann *et al*. (2018a) (*n* = 12), Krause-Kyora *et al*. (2018) (*n* = 3), Barquera *et al*. (2020) (*n* = 1), Patterson Ross *et al*. (2018) (*n* = 1), Kahila Bar-Gal *et al*. (2012) (*n* = 1) and Neukamm *et al*. (2020) (*n* = 1). We further included ancient genomes with high coverage from Eastern Eurasia (Sun *et al*., 2024) (*n* = 25), one additional ancient genome from Mexico (HSJN194, 1472–1625 CE) (Guzmán-Solís *et al*., 2021) and 264 modern genomes (sampling time range: 1973-2013). The modern genomes selection is to a large extent based on the complete genome data analyzed by Harrison *et al*. (2011), who studied the impact of the Hepatitis B e-Antigen (HBeAg) status on viral evolutionary rates, and includes genotype A, C, D, E and F genomes. To assess the plausibility of previously suggested associations of HBV divergence with human migration events, this selection was further augmented with four genotype H genomes, 55 genotype B genomes from Bouckaert *et al*. (2017) and 21 genotype A3 genomes from Andernach *et al*. (2009). To obtain a broad representation of HBV genotypes, we incorporated a further four genotype G genomes and one putative genotype J genome. One modern genome (of genotype A) was newly obtained here from a serum sample from the DRC dating back between May 1988 and 1990 (cfr. Supplementary Information). Finally, the modern HBV genomes also include twelve genomes obtained from non-human primates previously also included in the analyses of Mühlemann *et al*. (2018a) and Kocher *et al*. (2021).

We aligned the total of 414 genomes using MAFFT 7.453 (Katoh *et al*., 2009) and applied the same masking of ambiguously aligned regions as in Kocher *et al*. (2021). HBeAg status or predicted status was retrieved from the literature or inferred based on the precore G1896A stop codon mutation Harrison *et al*. (2011). Sample ages were set to the time in years since 2013 (the most recent sampling year in the genome data set).

### Recombination analyses and temporal signal analysis

We used RDP5.58 (Martin *et al*., 2021) to detect and characterize individual recombination events that occurred during the evolution of the HBV genomes represented in the full data set (*n* = 414). Default program settings were used throughout. Briefly, a fully exploratory automated scan with the RDP (Martin *et al*., 2021), GENECONV (Sawyer, 1989), and MaxChi (Smith, 1992) methods was initially used to detect recombination signals (these methods were used in ‘primary scanning mode’), and the Bootscan (Martin *et al*., 2021), Chimaera (Posada and Crandall, 2001), SiScan (Gibbs *et al*., 2000), and 3Seq (Lam *et al*., 2018) methods were used to verify these signals (these latter four methods were used in ‘secondary scanning mode’). Then RDP5.58 verified whether each detected recombination event impacted the topology of the HBV phylogenetic trees, flagging phylogenetically impactful recombination events for later removal from the alignment. To remove individual recombination events, RDP5.58 then identified recombinant sequences using a group of recombinant identification tests based on the EEEP (Beiko and Hamilton, 2006), PHYLPRO (Weiller, 1998) and VISRD (Lemey *et al*., 2009) methods. With the likely locations of recombination breakpoints and the identities of recombinant sequences in hand, the tracts of sequence bounded by the break-points of all flagged phylogenetically impactful recombination events were removed from all identified recombinant se-quences by replacing the tracts of recombinationally derived sequence with gap characters. Similarity plotting using a sliding window approach was performed using SimPloT++ (Samson *et al*., 2022) with a window size of 750 years BP, a step size of 30 and an HKY substitution model with gamma-distributed among-site rate heterogeneity.

We used IQ-TREE 2.1.2 (Minh *et al*., 2020) to reconstruct maximum-likelihood phylogenetic trees under a general time-reversible (GTR) model with a ‘FreeRate’ model of among-site rate heterogeneity (Soubrier *et al*., 2012). Root-to-tip divergences were extracted from these phylogenies and plotted against sampling time using TempEst (Rambaut *et al*., 2016).

### Bayesian time-measured phylogenetic inference

#### HBV molecular clock modeling

We build on previous efforts in mixed-effects molecular clock modeling (Bletsa *et al*., 2019; Vrancken *et al*., 2014) and TDR modeling (Membrebe *et al*., 2019) to develop a molecular clock model capable of accommodating different sources of HBV evolutionary rate variation. Specifically, we adopt a branch-specific modeling approach that combines both fixed- and random-effects, and that follows the general formulation of Bletsa *et al*. (2019) for the substitution rate parameter *µ*_*i*_ on branch *i*:

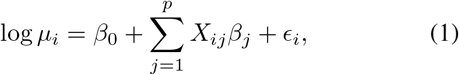

where *β*_0_ is the unknown intercept, *β*_*j*_ is the fixed-effect size of the *j*th covariate *X*_*ij*_ (out of *p* covariates), and _*i*_ are independent and normally distributed random-effect variables with mean 0 and an estimable variance. To accommodate time-dependency as a fixed-effect on the rates, we consider a covariate as the time at the midpoint of branch *i*. This covariate therefore does not represent strictly fixed values, but estimable parameters in our phylogenetic model. In our HBV application, we also consider two fixed-effects that follow standard specifications in the mixed-effects model (Bletsa *et al*., 2019). We allow for a potential host effect on the rate by setting the *X*_*i*_ = 1 for branches in a non-human primate virus lineage and *X*_*i*_ = 0 for all other branches (in human virus lineages). To model the potential effect of the HBeAg status on the rate along the tip branches leading to the sampled genomes, we set *X*_*i*_ = 1 for tip branches representing genomes associated with an HBeAg negative status and *X*_*i*_ = 0 for tip branches representing genomes associated with an HBeAg positive status. In our Bayesian inference, we specify a normal prior with a mean of -10 and a standard deviation of 10 on *β*_0_ and a normal prior with a mean of 0 and a standard deviation of 2 on the *β*_*j*_’s. For the TDR effect, we also consider a prior specification based on the posterior estimate for the TDR coefficient in a lentivirus data set analyzed by Membrebe *et al*. (2019) (𝒩 (*µ* =*™* 0.621, *σ* = 0.035)).

Our mixed-effects model considers time dependency for the rates over all branches in the phylogeny. However, recent mechanistic modeling of time-dependent rates due to substitution saturation argued that, going back in time, such saturation only starts to take effect at some time since the present (Ghafari *et al*., 2021). To model this behaviour, we also extend the fixed-effects specification for time-dependent rates such that the effect only applies at a time older than an estimable threshold since the present (cfr. Supplementary Information). We implement the mixed-effects model with branch-specific time-dependent rate variation (TDRmix), as well as its threshold extension, in BEAST X (Baele *et al*., 2025b). Initial inference attempts using vague prior specifications indicated challenges to obtain an informed estimate for the threshold parameter. Therefore, we specified a normal prior with a mean of 187.5 years and a standard deviation of 60 years based on the estimate for group VII RT-DNA viruses by Ghafari *et al*. (2021).

#### Hamiltonian Monte Carlo inference

In relaxed molecular clock formulations, relatively strong correlation between branch-specific rates and node heights may hamper efficient MCMC inference. This is further exacerbated by parameterising the branch-specific rates as a function of unknown midpoint times of the branches in our TDRmix model. Not surprisingly, initial explorations using standard random walk transition kernels to sample from the model posterior distribution demonstrated serious mixing problems. We therefore confront this problem by applying a Hamiltonian Monte Carlo (HMC) method to simultaneously sample all the branch-specific rates from the model posterior distribution (Ji *et al*., 2020). HMC is a state-of-the-art Markov chain Monte Carlo (MCMC) method that produces distant proposals with relatively high acceptance rate for the Metropolis algorithm by exploiting numerical solutions of the Hamiltonian dynamics (Neal *et al*., 2011). We employ HMC transition kernel implementations for the branch-specific rates and, following the ratio transformation approach by (Ji *et al*., 2023), for the node heights as well as for the rates and node heights simultaneously. To make HMC computationally tractable for large data sets, we employ the linear-time gradient algorithm implemented in the BEAGLE library (Ayres *et al*., 2019; Gangavarapu *et al*., 2024) with access in BEAST X (Baele *et al*., 2025b).

#### Other model and inference specifications

In our full probabilistic Bayesian inference, we partition the genome alignment into overlapping and non-overlapping gene regions and allow for different relative substitution rates between the alignment partitions. We specify a GTR substitution model and allow for separate discretized gamma-distributed among-site rate variation in both the overlapping and non-overlapping gene region partitions. We specify a flexible non-parametric skygrid coalescent prior on the tree (Gill *et al*., 2013) and employ recently developed HMC sampling to estimate the population size parameters (Baele *et al*.,2020). We incorporated sampling time uncertainty by specifying prior distributions over the ages of the ancient genomes and estimating their age as part of the MCMC inference. If age intervals were associated with genomes based on radiocarbon dating in the original studies, we specified a normal prior with a mean set to the midpoint of the interval, shifted relative to 2013 (time 0 in our analyses), and standard deviation such that 95% of the prior mass covers the age-interval. When age intervals were not provided but radiocarbon dating standard deviations were available, we estimated the standard deviation for the normal prior using a linear regression model. The model was trained on samples with known age intervals, using their calculated standard deviations as the response variable and their mean ages and radiocarbon standard deviations as predictors. When only a mean age was available, we used a prediction based on that age as the sole predictor. When age intervals were available based on other data, we used uniform prior specifications based on the shifted interval boundaries. For one viral sample, for which only a mean age was available, the uniform interval boundaries were predicted from its age based on the relationship between age and the boundaries for the samples that had this available. Because there was no sampling time available for three modern genotype G genomes, we specified an exponential prior on their age with a mean of two and an offset set to the submission date of the genomes. For the newly obtained modern genotype A genome from the DRC, we set a two-year uniform prior on its age based on a sampling date between 1988 and 1990. In all analyses using different molecular clock models, we constrained the basal lineage comprising genotype F and H to be monophyletic. XML input files for BEAST are available at https://github.com/phylogeography/HBV_TDR. We diagnose MCMC runs using Tracer 1.7.3 (Rambaut *et al*., 2018) and ensure that effective sample sizes are larger than 100 for all continuous model parameters, often achieved by combining several independent runs. We obtain highest independent posterior subtree (HIPSTR) summary trees using TreeAnnotator X (Baele *et al*., 2025a).

### Phylogeographic reconstructions in continuous space

We aim to infer the historical spread of HBV across Eurasia, but the sampling, and importantly, available migration-based calibrations, extend beyond Eurasia. We therefore take a two-step procedure and first infer trees scaled in time units by fitting a mixed-effects molecular clock model to the full data set excluding the primate sequences and the putative genotype J genome (AB486012) that descends from the Asian primate lineage. Next, we fit a spatial diffusion model to the resulting set of posterior trees, pruned to lineages of interest and only including ancient samples to avoid the impact of modern migration. In the first step, we employ the threshold extension with an informative prior, and combine dated tips, the informative prior on the TDR effect and three known human migrations as calibration information in a ‘total-evidence’ approach. For the migration-based calibrations, we use prior distributions incorporating migration time uncertainty and a novel approach to integrate over all possible times for migration events along a specific branch. This avoids having to rely on the often made assumption that the migration time applies to the node height of a particular clade. We achieve this by introducing an additional tree node with estimable height along the branch ancestral to the clade of interest. As prior distributions for the height of this node, we use a lognormal distribution for the trans-Atlantic slave trade with a mean of 4.45, a standard deviation of 0.65 and an offset of 150 years (based on the number of captives disembarked per year from https://www.slavevoyages.org), and uniform priors for the movement of Neo-Eskimo (Inuit) populations into the circumpolar Arctic (647-953 years BP; Bouckaert *et al*. (2017)) and the early migration to the Americas via the Beringia land bridge (18 000-25 000 years BP, based on ancient genomics: Llamas *et al*. (2016); Moreno-Mayar *et al*. (2018); Raghavan *et al*. (2015)). For the latter scenario, we also used a uniform prior on the corresponding height of the node representing the relevant clade (the genotype F and H tMRCA) (14 600-17 500 also based on ancient genomics: Llamas *et al*. (2016); Moreno-Mayar *et al*. (2018); Raghavan *et al*. (2015)). The posterior tree distribution resulting from the total-evidence molecular clock analysis was pruned for subsequent phylogeographic inference of specific lineages, retaining only the ancient genomes belonging to these lineages. We use a relaxed random walk (RRW) model to infer dispersal patterns through time (Fisher *et al*., 2021; Lemey *et al*., 2010), while we adopt a BEAST implementation of the integrated Brownian motion (IBM) model to estimate dispersal rates (Bastide *et al*., 2024), which avoids sampling intensity bias (Dellicour *et al*., 2024). For both approaches, we perform analyses with diffuse priors on the root locations as well as with root locations constrained to specific areas while integrating over uncertainty in these areas. We use node-specific dispersal velocities estimated in the IBM model to summarize velocity trends by binning values into time intervals and computing their mean and standard deviation within each bin.

### Temporal signal evaluation

We evaluate temporal signal using a recently developed Bayesian model testing approach (BETS, Duchene *et al*. (2020)) as well as using date-randomization testing (Ramsden *et al*., 2008). We perform both tests using the same substitution model as in the previously described BEAST analyses, a strict molecular clock model and a constant population size model as tree prior. In the BETS approach, we adopt generalized stepping stone sampling (GSS; Baele *et al*. (2016)) to estimate (log) marginal likelihoods for a model with dated tips (and an estimable substitution rate) and a model with contemporaneous tips (and a fixed substitution rate). The GSS used a matching coalescent model as tree working prior and relied on 50 path steps sampling 1 million states each to sample from a series of power posteriors. The date-randomization test compares posterior rate distributions from dated tip models applied to the actual sampling dates with posterior rate distributions from an MCMC run in which the dates are shuffled using a randomization transition kernel in BEAST (Trovão *et al*., 2015).

### Simulations

To evaluate the performance of the branch-specific TDR model with random-effects, we performed two sets of simulations that are modeled after a subset of the HBV genome collections. Specifically, we used a subset (*n*=18) of ancient genomes and 17 modern genomes distributed across the HBV phylogeny. We analyzed this data set using an uncorrelated relaxed molecular clock model and a branch-specific TDR model with random-effects and used the parameter estimates for the subsequent simulations. We simulated 20 replicates under both an uncorrelated relaxed molecular clock model (Drummond *et al*., 2006) and an epoch TDR model (Membrebe *et al*., 2019). We subsequently analyzed both sets of simulations using the new branch-specific TDR model with random-effects.

## Results

### Various sources of HBV evolutionary rate variation

We explore sources of evolutionary rate variation in HBV genome data from both ancient and modern samples, primarily from humans, but also including a limited number of genomes from non-human primates. Upon recombination filtering (cfr. Methods), we visually examine temporal signal by plotting root-to-tip divergence as a function of sampling time (Figure 2). While this graphical exploration illustrates the accumulation of genetic divergence over a timescale of more than 10 000 years, there is substantial variation in this pattern. This is, for example, noticeable for the modern samples for which some degree of variation may be explained by differences between human and non-human primate lineages (purple primate tips in Figure 2), perhaps as the result of host-specific evolutionary rates. A generally higher root-to-tip divergence can be observed for ancient genomes associated with an HBeAg negative status (Figure 2B). A higher substitution rate for HBV genomes associated with an HBeAg negative status has previously also been demonstrated for modern samples, both by longitudinal sampling from HBV patients and by phylogenetic comparison across hosts (Harrison *et al*., 2011; Vrancken *et al*., 2017). Recent modeling work has explained this by higher rates of clearance of HBV covalently closed circular DNA (cccDNA) during the HBeAg negative status (Lythgoe *et al*., 2021).

**FIG. 2.**
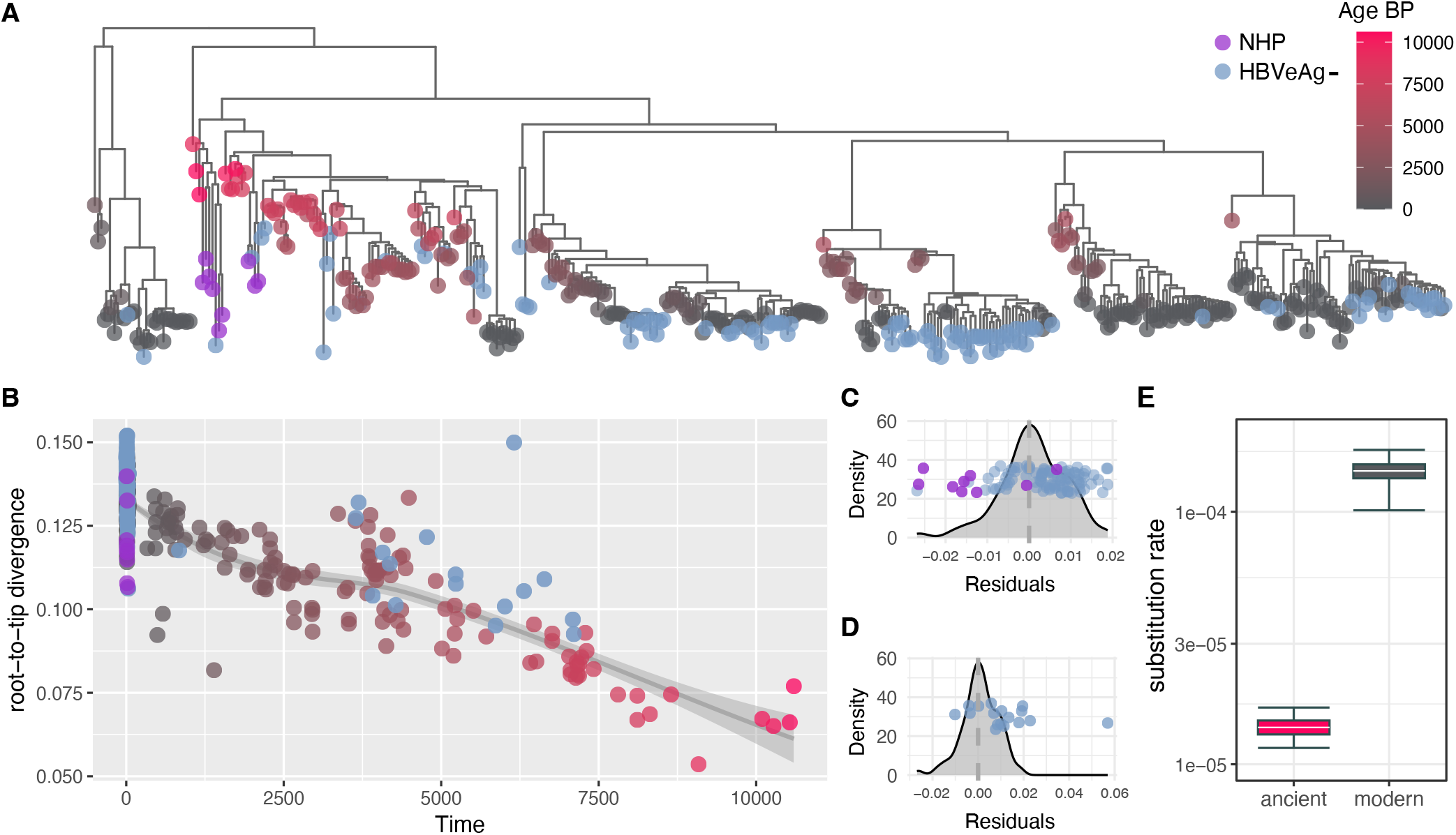
Maximum-likelihood phylogenetic reconstruction, root-to-tip divergence as a function of sampling time, its residuals, and evolutionary rate estimates for ancient and modern HBV genomes. (A) In the phylogeny, the red-to-gray tip circle color gradient reflects the sampling time or age BP of the genomes from human samples. Genomes with an HBeAg negative status are depicted with a blue tip color whereas genomes sampled from non-human primates (NHP) are depicted with a purple tip color. (B) The same tip symbols and colors are maintained in the root-to-tip divergence plot. The grey line and 95% confidence interval area represent a local (LOESS) regression through the genomes sampled from humans with an HBeAg positive status. (C) Residual distribution for root-to-tip divergences as a function of sampling times for the modern genomes. Residual data points are superimposed for HBV genomes from NHPs and HBeAg negative patients. (D) Residual distribution for root-to-tip divergences as a function of sampling times for the ancient genomes. Residual data points are superimposed for HBV genomes from HBeAg negative patients. (E) Evolutionary rate estimates under an uncorrelated relaxed molecular clock model fitted separately to the ancient (red-filled box) and modern human HBV genomes (gray-filled box). The box plots summarize the median estimate (white line), the interquartile range (box), and 95% HPDs (whiskers).

In order to assess whether the timescale of measurement also affects HBV evolutionary rates and patterns of divergence over time, we independently estimated evolutionary rates for both the modern and ancient human genomes using Bayesian inference under an uncorrelated relaxed molecular clock model (Figure 2C). We infer about a tenfold higher rate from modern genomes, which corroborates differences in estimates for similar ancient and modern data sets (Figure 1, Table I). Formal testing confirms that both the ancient and modern genomes contain sufficient temporal signal (Table I and Supplementary Figure 1). Taken together, these estimates underscore the need to account for time-dependent rate (TDR) effects when combining modern and ancient HBV genomes, consistent with a phenomenon documented across multiple viruses with diverse evolutionary timescales (Aiewsakun and Katzourakis, 2016; Duchêne *et al*., 2014a).

**Table I.** Evolutionary rate estimates and temporal signal tests for ancient and modern HBV genomes. Model support is quantified using Bayes factors (BFs), expressed on the natural log scale. Date randomization tests are in agreement with the outcomes of the BETS approach (Supplementary Figure 1).

| Data | sampling time range<br>(years before present) | mean substitution rate (HPD) $10^{-5}$ subst./site/yr | ln BF (dated vs.<br>contemporaneous) |
| --- | --- | --- | --- |
| Ancient & modern | 0 – 10 598 | 1.49 (1.31–1.68) | 715 |
| Ancient | 333 – 10 598 | 1.41 (1.16–1.68) | 338 |
| Modern | 0 – 40 | 18.5 (13.2–23.4) | 31 |

### HBV molecular clock modeling

We confront the vari-ous sources of HBV evolutionary rate variation through the development of a flexible molecular clock approach. Specifically, we implement a branch-specific version of a TDR model (Membrebe *et al*., 2019) in a Bayesian framework that allows combining TDR with host-specific rates and a potential impact of the HBeAg status on the tip branch rates. In addition, we also consider unexplained branch-rate variation through random-effects, which extends our fixed-effects molecular clock model into a mixed-effects model (Bletsa *et al*., 2019; Vrancken *et al*., 2014).

We demonstrate that the branch-specific version of the TDR model can accurately capture TDR dynamics simulated under an epoch structure while avoiding false positive TDR effect identification when simulations are performed under a uncorrelated relaxed clock model (Supplementary Figure 2). When fitting the branch-specific TDR molecular clock model with only fixed-effects to the HBV complete genome data set (Figure 3A), we estimate a significant negative rate effect (on a log scale) for the non-human primate lineages, a significant positive rate effect on the tip branches for genomes sampled during the HBeAg negative status of infection, and a significant time-dependent decay in evolutionary rate throughout the evolutionary history. When also considering random-effects on the branch-specific rates (Figure 3B), we recover somewhat more pronounced HBeAg and TDR effects, but now a positive effect on the rates for the non-human primate lineages with a large credible interval. This indicates that the HBeAg and TDR effects provide an improved fit for the mixed-effects model in comparison to the uncorrelated relaxed molecular clock model. The difference in non-human primate rate effect between the fixed-effects and mixed-effects model may be due to an identifiability issue in the mixed-effects model because the lower divergence for the two non-human primate lineages (Figure 2) may also be explained by a negative random-effect on the branches ancestral to these two lineages. The posterior mean estimate for the standard deviation of the normally distributed random variables _*i*_ (0.50, 95% highest posterior density (HPD) interval: [0.44,0.57]) in the mixed-effects clock model indicates substantial rate variation in addition to the fixed-effects. Based on these estimates, we reduce the mixed-effects model to only include the HBeAg and TDR effects together with the random-effects, but we now consider a model extension that allows TDR to take effect after some estimable time since the present (Figure 3C). This is inspired by recent developments that model TDR as a function of sequence saturation (Ghafari *et al*., 2021), which is expected to begin at some time point in the past. Using an informative prior for this time threshold (cfr. Methods), we estimate a posterior mean threshold of 43 years (95% HPD: [0.23,89]) indicating that the evolutionary rate starts to decline roughly after the modern sampling of HBV. This has little impact on the posterior estimates of the two fixed-effects (Figure 3C). The negative TDR effect implies a substantial rate decline into the past, but the effect estimates are somewhat lower than those previously obtained using a similar model (Membrebe *et al*., 2019) (Figure 3A-C). It is plausible that the different sources of rate variation confound an accurate estimation of the TDR effect on the timescale of HBV evolution. Therefore, we also fit the same model with an informative prior on the TDR effect based on previous estimates (cfr. Methods), which results in a more pronounced and more precise TDR estimate as expected (Figure 3D). The posterior mean estimate for the intercept in this model (*β*_0_) reflects an evolutionary rate of 5.4 *×*10^*™*4^ (95% HPD: [3.9 *×*10^*™*4^, 7.6 *×*10^*™*4^]) subst./site/yr whereas the rate estimate on the branch with the deepest midpoint is at least two orders of magnitude lower (3.0 *×*10^*™*6^; 95% HPD: [7.4 *×*10^*™*7^, 6.2 *×*10^*™*6^] subst./site/yr).

**FIG. 3.**
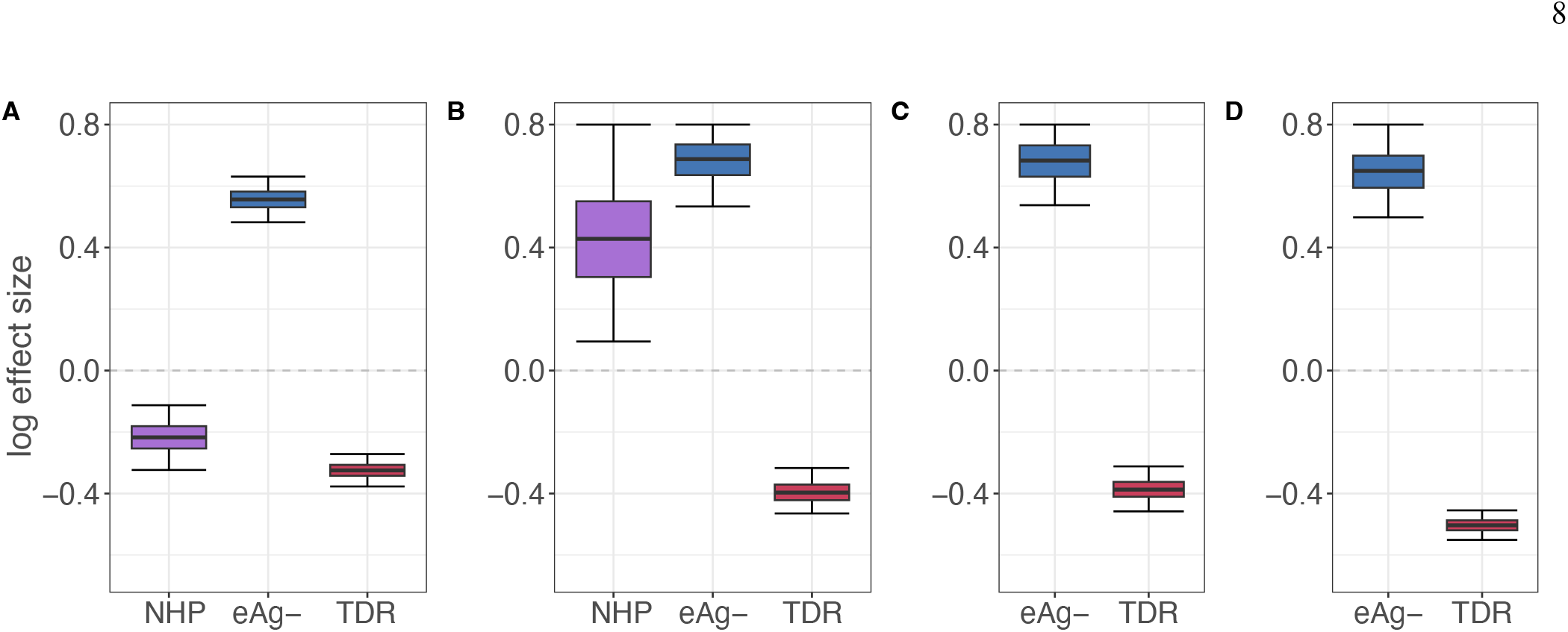
Molecular clock model estimates. Marginal posterior distributions for the log fixed-effect sizes in the molecular clock models. Estimates are shown for a fixed-effects model (A), a mixed-effects model (B), a mixed-effects model with a time threshold on the TDR effect (C) and a mixed-effects model with a time threshold and an informative prior on the TDR effect (D). As fixed-effects we consider an evolutionary rate effect on the branches associated with the non-human primate lineages (‘NHP’, in (A) and (B)), an evolutionary rate effect on the tip branches for genomes sampled during the HBeAg negative state of human infection (A-D), and a time-dependent rate (TDR) effect on all branches as a function of their unknown midpoint times (A-D). The box plots summarize the median estimate (black line), the interquartile range (box), and 95% HPDs (whiskers).

### Reconciling HBV diversification with historical human migration

The strong rate decline we infer under the TDRmix model has a specific impact on divergence time estimates. Relative to estimates under time-homogeneous molecular clock models, deep divergence times are pushed further towards the past whereas recent divergent times are pulled towards the present. We evaluate how this TDR effect shapes divergence times for events that have been previously associated with human migration (Figure 4).

**FIG. 4.**
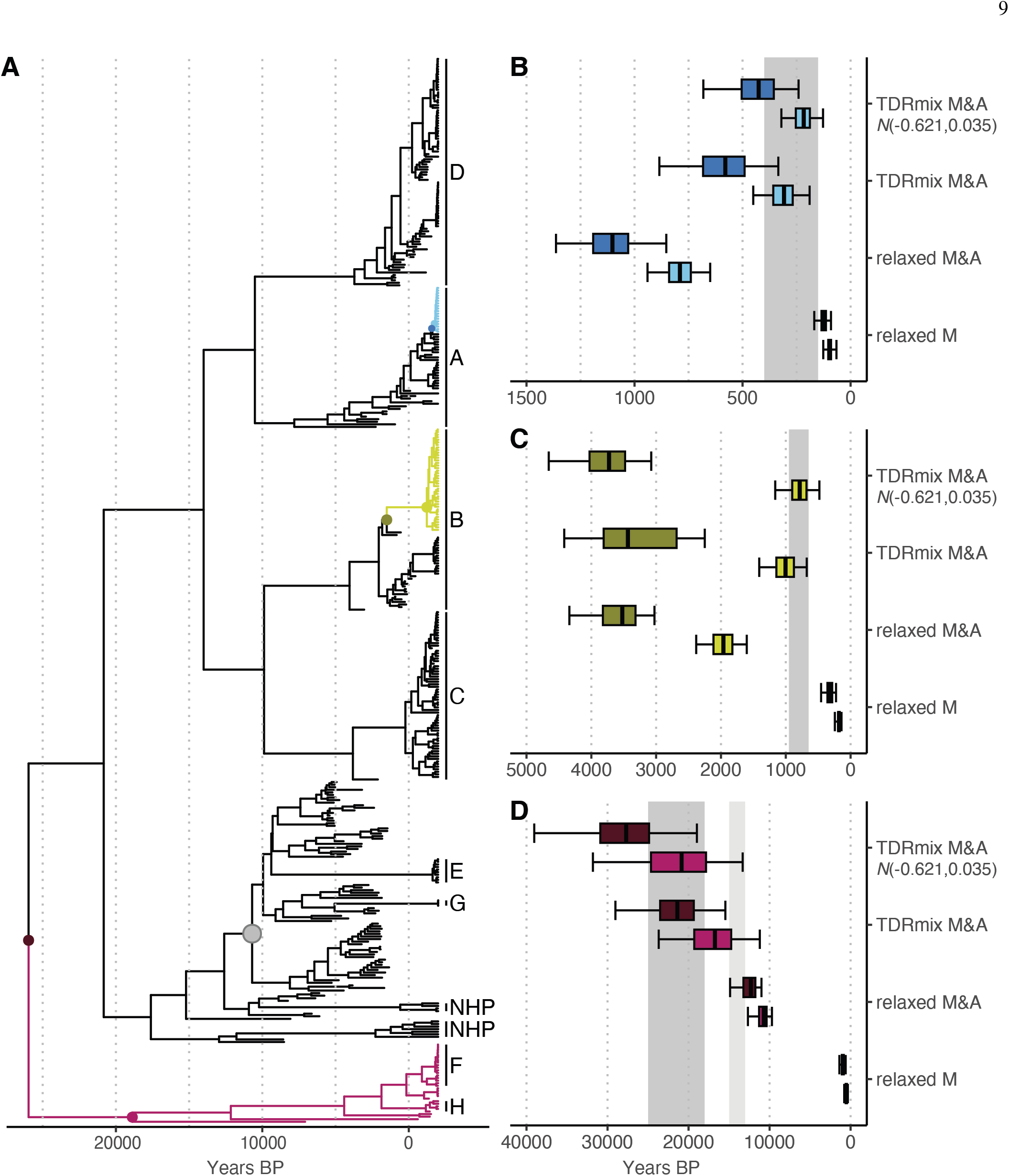
HBV divergence time estimates and historical human migrations. tMRCA estimates are summarized for three specific clades in the time-measured phylogeny (panel A). The three clades represent the (quasi-)subgenotype A3 clade in Haiti (blue), subgenotype B5 circulating in the circumpolar Arctic (yellow-brown), and the basal genotype F/H lineage identified in Amerindian populations (purple). HBV genotype A-E and two non-human primate (NHP) lineages are indicated at the tips of the tree. In some instances, basal viral diversity from ancient samples are included in the genotypes for simplicity. For subgenotype B5 and lineage F/H, both the tMRCA nodes and their parent nodes are indicated with a filled circle. The Western-Eurasia Neolithic-to-Bronze-Age (WENBA) lineage is indicated with a grey filled circle. As divergence times for the (quasi-)subgenotype A3 clade in Haiti (B), subgenotype B5 (C), and the basal F/H lineage (D), both the estimates for the tMRCAs (lower box) as well as for their parent nodes (upper box) are plotted because migration events could have occurred along the branches connecting these nodes. We summarize estimates for the TDRmix (with vague and 𝒩 (*µ* =*™* 0.621, *σ* = 0.035) prior) and relaxed clock model applied to the full data set (modern and ancient: M&A) as well as for the relaxed clock model applied to modern genomes only. The box plots summarize the posterior median (black line), interquartile range (box), and 95% HPDs (whiskers). The historical human migrations that have been associated with the three lineages are indicated with grey rectangles: (B) the transatlantic slave, (C) the movement of Inuits into the the circumpolar Arctic between 647-953 years BP and (D) the first colonization of the Americas between *∼* 25 000 and 18 000 years BP (and also the Bølling-Allerød interstadial from *∼* 15 000 to 13 000 years BP is depicted in lighter grey).

One of these events is the divergence of the Amerindian genotypes F and H that together form a basal lineage in the HBV phylogeny (Figure 4D), which has been associated with early human migrations responsible for the peopling of the Americas (Paraskevis *et al*., 2013). Widespread human expansion took place during the Bølling-Allerød interstadial in North America, a warm interstadial period that covered the final stages of the Last Glacial Period from *∼* 15 000 to 13 000 years BP (Palacios *et al*., 2020; Waters, 2019). However, ancient human DNA studies have provided evidence that the ancestors of the First Americans started to diverge from Eurasian relatives between *∼* 25 000 and 18 000 years BP (Llamas *et al*.,2016; Moreno-Mayar *et al*., 2018; Raghavan *et al*., 2015). As such migration events could have occurred at any point along the branch leading to the MRCA of the basal F/H lineage, we also consider time estimates for the parent node from which the lineage diverged. We summarize estimates for the TDRmix and uncorrelated relaxed clock model, and also fit the latter to only modern genomes for comparison (Figure 4D). As previously demonstrated by Mühlemann *et al*. (2018a), relaxed clock model estimates for the combined modern-ancient data are largely inconsistent with the first colonization of the Americas as only the posterior density for the parent node of the genotype F/H MRCA (13 385; 95% HPD: [12 179-14 801]years BP) overlaps with the Bølling-Allerød interstadial but not with the early divergence between *∼* 25 000 and 18 000 years BP (Figure 4D). In contrast, the TDRmix model yields a compatible estimate with this early divergence, in particular when the informative prior on the time-dependent decline rate is used.

Next, we focus on HBV subgenotype B5, which is endemic in indigenous populations of the circumpolar Arctic. Its introduction in this area has been associated with the movement of Neo-Eskimo (Inuit) populations between 647-953 years BP (Bouckaert *et al*., 2017). While divergence time estimates for the combined modern and ancient data are now too old under the relaxed clock model to be compatible with this migration, the posterior age densities for subgenotype B5 MRCA under the TDRmix model overlap with the migration time range (Figure 4C), 1 004 (95% HPDs: [675-1 410]) and 785 (95% HPDs: [482-1 164]) for a vague and informative prior on the TDR decline rate respectively).

Finally, we consider (quasi-)subgenotype A3 in Haiti, a clade that branches off from Nigerian strains and that was presumably introduced during the trans-Atlantic slave trade (Andernach *et al*., 2009; Paraskevis *et al*., 2013). Also in this case, divergence time estimates under a relaxed clock model for the combined modern and ancient data are too old to fit with this hypothesis, but the TDRmix model makes the Haiti A3 tMRCA estimate overlap with the time interval for the slave trade, in particular when the informative prior on the TDR decline rate is used (Figure 4C).

### A ‘total-evidence’ spatiotemporal reconstruction captures historical human migrations

Motivated by the concordance between viral divergence times and various human migration events across different timescales, we further unravel the HBV spatiotemporal spread using a ‘total-evidence’ approach. We focus only on human HBV genomes and combine dated tips, calibration information for the three studied migration events in the TDRmix model and prior information on the TDR effect (cfr. Methods). Using continuous diffusion models for spatially explicit phylogeographic reconstructions, we first focus on the Western-Eurasia Neolithic-to-Bronze-Age (WENBA) lineage (Figure 4A), which includes numerous genomes recovered from early European farmers (Kocher *et al*., 2021). Whether this lineage originated in Near Eastern centers of early agriculture or in another location along early expansion routes has so far remained elusive (Kocher *et al*., 2021). This is challenging to test using ancestral reconstruction because of large estimation uncertainty and a predominant sampling of European WENBA genomes. However, relatively precise estimates for the speed of farming expansion have been drawn from archeological records, dating back to seminal work by Ammerman and Cavalli-Sforza (1971). Using radiocarbon-dated sites, the speed of the farming expansion across Europe was estimated at an average rate of approximately 1 km per year with an origin inferred in Jericho (Ammerman and Cavalli-Sforza, 1971). To test whether HBV spread followed the expansion of early farmers from Anatolia, we estimate the phylogeographic dispersal velocity of the WENBA lineage. Because summaries of dispersal velocity from widely-used continuous diffusion models are sampling-inconsistent (Dellicour *et al*., 2024), we adopt a novel integrated Brownian motion (IBM) model that allows accurate estimation of dispersal velocity (Bastide *et al*., 2024). We estimate a posterior mean HBV dispersal velocity of 1.05 (95% HPDs: [0.55,1.50]) km/yr in near perfect agreement with the archeological estimates of (Ammerman and Cavalli-Sforza, 1971) as well as more recent estimates based on additional data (Pinhasi *et al*., 2005) (Figure 5A). We obtain highly similar estimates (1.10, 95% HPDs: [0.69,1.57] km/yr) when constraining the WENBA root in the Near East (Figure 5A). The dispersal rate remained roughly constant throughout the WENBA evolutionary history (Figure 5B), in line with a wave-of-advance model (Ammerman and Cavalli-Sforza, 1971).

**FIG. 5.**
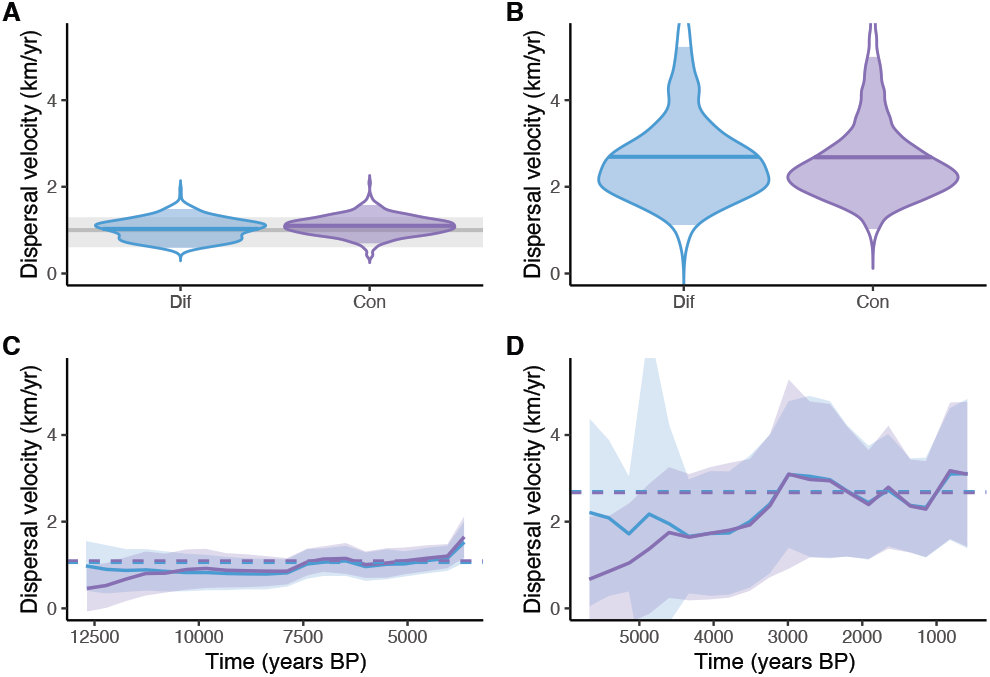
Dispersal rate estimates for the WENBA and genotype D lineage. Estimates were obtained using an IBM model with diffuse (Dif, blue) root location priors or constrained (Con, purple) root locations. (A) Posterior distributions for the WENBA mean dispersal rate. The horizontal grey line and ribbon represent archaeological estimates for the Neolithic migration rate (Ammerman and Cavalli-Sforza, 1971). (B) Posterior distributions for the genotype D mean dispersal rate. (C) Dispersal rate through time for the WENBA lineage. The solid line represents the mean dispersal rates across 50 time bins while the ribbon represents the standard deviations. The dashed lines represent the posterior mean estimate from (A). (D) Dispersal rate through time for the genotype D lineage – same is in (C) with posterior mean estimate from (B).

The RRW spatiotemporal reconstruction indicates that in the period up to 9 000 years BP (Figure 6B), the WENBA lineage had spread from Anatolia into the Balkans, initiating migration both along the Mediterranean/Cardial route and the Danubian route. In addition, north-eastward migrations from the Balkans may have contributed to the Cucuteni–Trypillia populations in present-day Romania, Moldova, and Ukraine, which carried a strong Anatolian farmer genetic component (Immel *et al*., 2020; Mathieson *et al*., 2018), and from which we infer further dispersal (Figure 6C). We reconstruct subsequent dispersal from this Eastern European area and from the Balkans into Central Europe (Germany, Figure 6B&C) in line with Central European Linear Pottery culture (Linearbandkeramik) farmers having predominantly Anatolian Neolithic ancestry (Haak *et al*., 2010; Mathieson *et al*., 2015). Considerable WENBA ancestry also reflects migration along the Cardial ware or maritime route. Still in the early Neolithic (Figure 6C), this results in expansion into Western and Southern Europe, including independent introductions into the Mediterranean islands of Corsica and Sicily. In the Copper and Bronze Age (Figure 6D), we infer further spread to into Spain, and additional spread to Southern Italy and Mediterranean islands. Bronze Age dispersals also include movements from Central Europe (Germany) to Scandinavia (Sweden) and from Russia to China (Figure 4A). Not only do our reconstructions corroborate established patterns of Neolithic migration, but they also provide fine-grained spatiotemporal insights that enhance our understanding of prehistoric population dynamics.

**FIG. 6.**
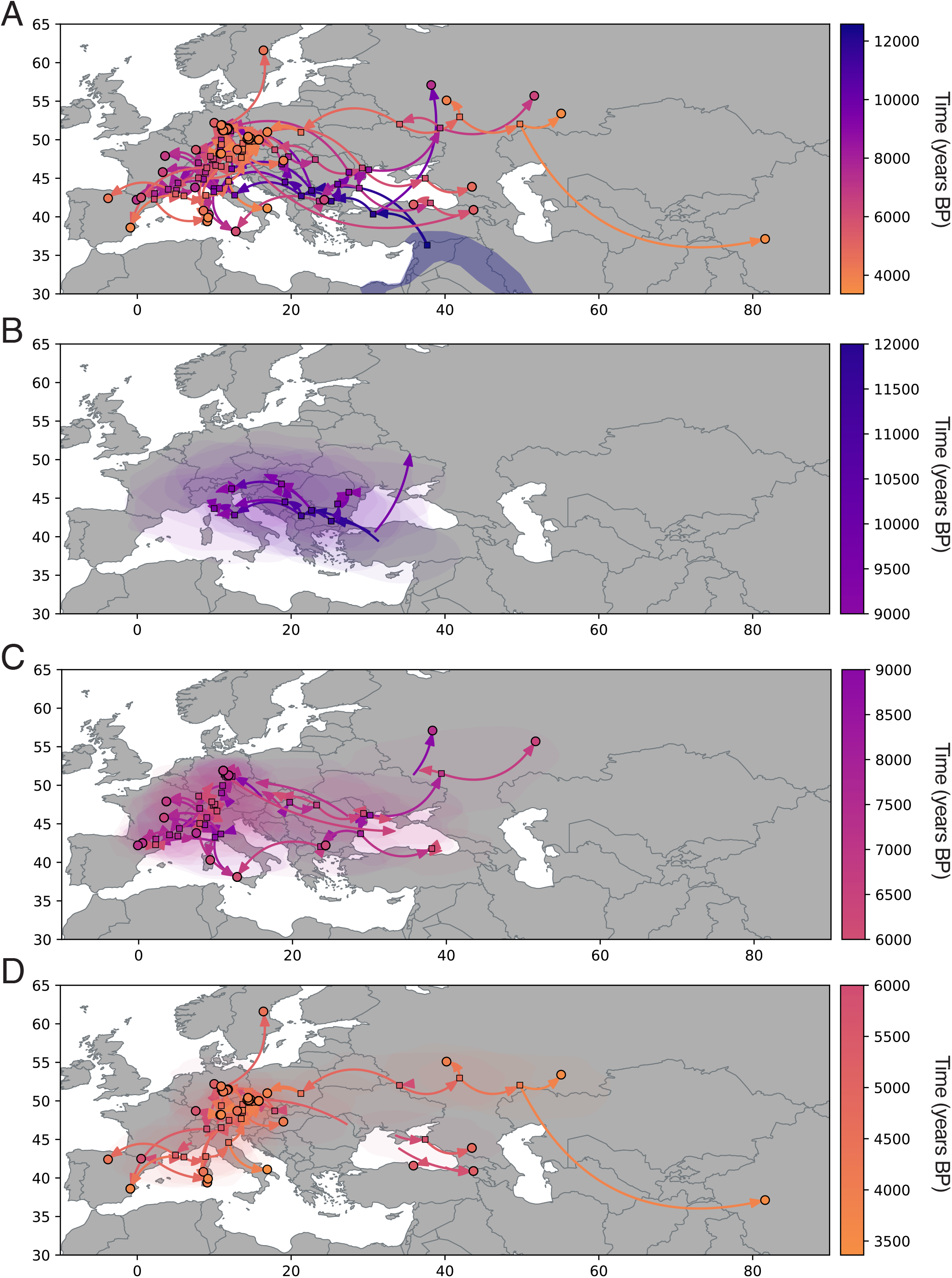
Reconstruction of the spatiotemporal spread of the Western-Eurasia Neolithic-to-Bronze-Age (WENBA) lineage. (A) Spatial projection of the highest independent posterior subtree for the relaxed random walk analysis of the complete WENBA lineage with a root constrained to the Near East. Internal nodes are represented by squares, while tips are represented by circles; arrows depict branches. The blue polygon that contains the root node represents part of the fertile Fertile Crescent. (B) Spatial projection up to 9 000 years BP (as in A). Arrows may represent partial branches if they extend beyond 9 000 years BP. Transparent polygons shown in this and subsequent panels represent the 80% HPD regions of the internal node locations. (C) Spatial projection from 9 000 to 6 000 years BP (as in B). (D) Spatial projection from 6 000 years BP to the most recently sampled ancient WENBA genome (as in B).

Apart from genotype G and E descendants (Figure 4), the WENBA lineage became extinct, and current HBV genotype distributions indicate that genotype D dominates across Eurasia (Castaneda *et al*., 2021; Velkov *et al*., 2018). This prompts an inquiry into the factors that may have facilitated the emergence and establishment of HBV genotype D. A second migration that profoundly influenced European ancestry began around 5 000 years BP, involving the Yamnaya people from the Pontic-Caspian Steppe spanning parts of present-day Russia and Ukraine (Haak *et al*., 2015; Saag and Metspalu, 2025). The expansion of steppe-related ancestry dramatically altered the cultural and genetic landscape in Europe. To test whether HBV genotype D may have spread in association with this migration, we performed phylogeographic analyses of the ancient genotype D samples. The RRW reconstruction places the posterior mean root location within the Pontic-Caspian Steppe, albeit with considerable uncertainty (Supplementary Figure 3). Additionally, the genotype D tMRCA estimate (5 803 years BP; 95% HPD: 4 632–6 945) is consistent with the steppe migration scenario. In contrast to the more asymmetrical Neolithic migration, we reconstruct a more bidirectional (westward and eastward) migration from the Pontic-Caspian Steppe (Figure 7). This is in agreement with the pan-Eurasian footprint of the Yamnaya expansion (Haak *et al*., 2015; Narasimhan *et al*., 2019). The eastward expansion may, for example, correspond to the introduction of steppe ancestry into the Altai region, but our inferred timing appears too recent to coincide with the emergence of the Afanasievo culture (Allentoft *et al*., 2015; de Barros Damgaard *et al*., 2018). The westward spread into Europe appears to have followed primarily northern routes, with less ancestry moving into southern and southwestern regions compared to the WENBA spread (Figure 7). This pattern is consistent with a well-established signal in the European archaeogenetic record: a north–south cline in steppe and Anatolian ancestry proportions (Lazaridis *et al*., 2025). Moreover, the reconstructed patterns of HBV spread do not support a single large population migration wave, as different locations are reached via largely independent routes. This corroborates predictions from spatially explicit paleogenomic simulations that support a scenario of continuous human gene flow, involving both short-range dispersal and rare long-distance movements from eastern populations (Rio *et al*., 2021). Our phylogeographic IBM approach estimates a dispersal rate for genotype D that is more than twofold higher (posterior mean 2.68 km/yr; 95% HPD: 1.02–5.00) than that inferred for WENBA, broadly consistent with genetic estimates of migration-front speeds (Racimo *et al*., 2020). The genotype D dispersal rate appears to have been higher since about 3 000 years BP (Figure 5B), associated with more recent migrations than the initial Yamnaya expansion (Figure 7B&C).

**FIG. 7.**
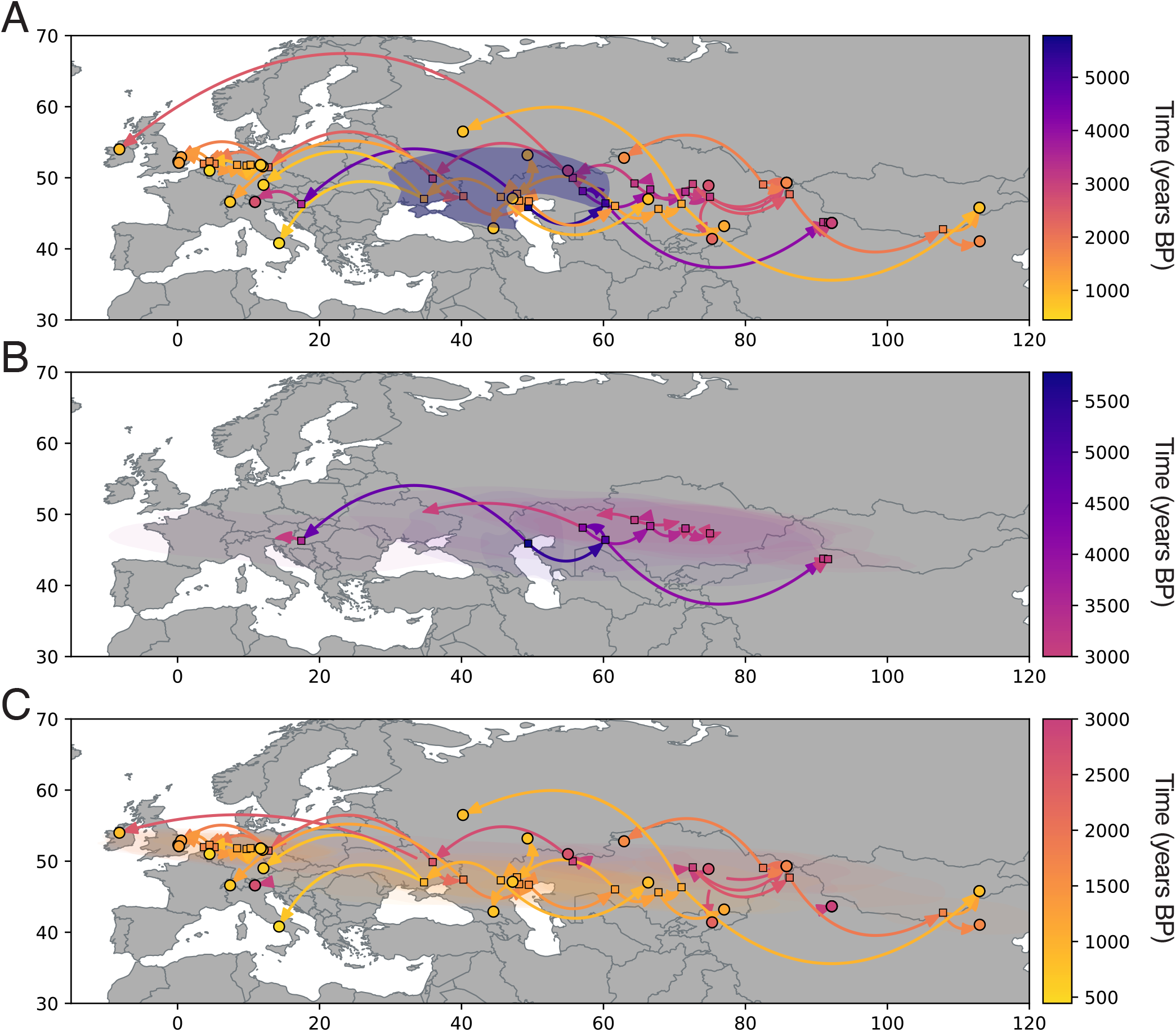
Reconstruction of the spatiotemporal spread of HBV genotype D. (A) Spatial projection of the highest independent posterior subtree for the relaxed random walk analysis of genotype D with a root constrained to the Pontic-Caspian Steppe. Internal nodes are represented by squares, while tips are represented by circles; arrows depict branches. The blue polygon that contains the root node represents the Pontic-Caspian Steppe area. (B) Spatial projection up to 3 000 years BP (as in A). Arrows may represent partial branches if they extend beyond 3 000 BP. Transparent polygons shown in this and the subsequent panel represent the 80% HPD regions of the internal node locations. (C) Spatial projection from 3 000 years BP to the most recently sampled ancient genotype D genome (as in B).

Multiple waves of migration and cultural interaction over millennia have resulted in a modern European genetic landscape characterized by admixture between Neolithic farmers and steppe pastoralists (Chintalapati *et al*., 2022). Despite the age lag between the WENBA (7 419-3 363 years BP) and ancient genotype D samples (2 958-444 years BP), our reconstructions indicate that their ancestry overlapped both in terms of population size trajectory and phylogeographic history (Figure 8). For instance, at 4 250 years BP the 95% Bayesian credible area for both lineages largely overlaps, in particular in Central Europe (Figure 8). This may have facilitated co-infections and, consequently, opportunities for HBV recombination. Evidence in support of this comes from genotype E, for which recombination has recently been reported (Kocher *et al*., 2021). Our analyses identified all genotype E genomes as recombinants that share a minor 728 bp recombinant segment defined by breakpoints in the polymerase–X overlap region and the core gene (cfr. Supplementary Information). Upon exclusion of this recombinant segment, genotype E clustered within the WENBA diversity (Figure 4). This observation is further supported by a similarity plot and by a phylogenetic reconstruction including only the minor recombinant segment of genotype E (Figure 8), both of which indicate an ancient genotype D ancestry. These findings illustrate how the spread of HBV through human migrations facilitated the emergence of novel viral diversity.

**FIG. 8.**
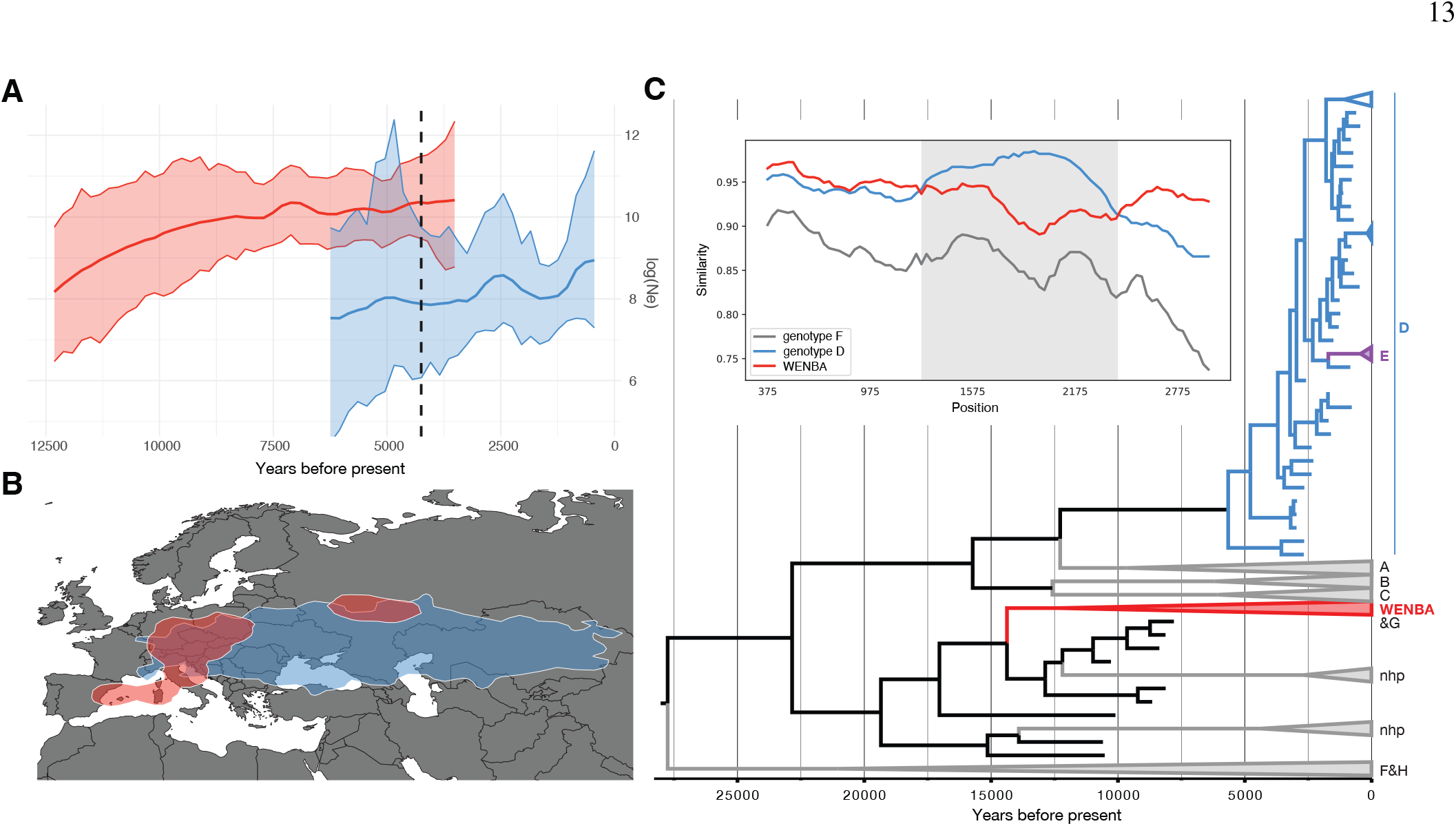
Genotype E as a recombinant of WENBA and genotype D. (A) The trajectory of log effective population size (Ne) through time for both the WENBA lineage and the ancient genotype D genomes. The vertical dashed line marks the time point (4 250 years BP) at which a phylogeographic summary was obtained for both lineages. (B) Summary of the 95% HPD region of uncertainty for the WENBA and ancient genotype lineages circulating at 4 250 years BP. For WENBA, the contour splits into two paths. (C) The time-calibrated phylogeny on the right represents the reconstruction under the TDRmix model including only the minor recombinant part of the genotype E genomes. Several lineages are collapsed to improve clarity, including the basal genotype F&H genomes (and related ancient samples), the non-human-primate lineages, WENBA, genotype A, B and C, genotype E and the two modern genotype D clades. The inset depicts a similarity plot with genotype E genomes as query and genotype D, WENBA and genotype F genomes as reference genomes. The minor recombinant region is highlighted in gray.

## Discussion

Estimating evolutionary rates and divergence times for HBV is plagued by extensive rate variation. In this study, we demonstrate that critical sources of evolutionary rate variation can be formally modeled in a molecular clock framework. Among the different effects we consider in our mixed-effects clock model, we estimate the largest effect sizes for the HBeAg state of human infection. A higher rate for HBV genomes associated with an HBeAg negative status has previ-ously been demonstrated by molecular studies (Harrison *et al*., 2011; Vrancken *et al*., 2017), but it has only been recently explained by higher rates of clearance of HBV cccDNA in hepatocytes (Lythgoe *et al*., 2021). Such cccDNA is converted from partially double-stranded relaxed circular DNA imported to the nucleus and serves as the transcriptional template for pregenomic and subgenomic RNAs. The relatively long life-span of cccDNA has been put forward as a simple explanation for the low HBV evolutionary rate compared to other RNA viruses with similar mutation processes (Lythgoe*et al*., 2017). Higher rates of cccDNA clearance during the HBeAg negative status result in a shorter lifespan of cccDNA, and hence a shorter viral generation time, which results in higher evolutionary rates (Lythgoe *et al*., 2021). We can only model the HBeAg status on tip branches for sampled viruses and therefore we are ignoring potential HBeAg status variability in the ancestral infection history. However, data from human transmission and small animal model studies imply that HBeAg-negative virus is much less likely to transmit and establish chronic infections (Kramvis, 2016; Tong and Revill, 2016; Zoulim *et al*., 1994), providing some reassurance that modeling tip branches largely captures the HBeAg effect.

The TDR effect we accommodate in our model is crucial in recovering divergence estimates that are compatible with human migrations both in the distant and more recent past. An important example of this is the divergence of the basal HBV lineage consisting of genotypes F and H, which has been associated with early migration to the Americas via the Beringia land bridge. Ancient genomics has demonstrated that this involved the movement of distinct and previously unknown populations from northeast Asia that diverged from their closest Eurasian relatives between *∼* 25 000 and 18 000 years BP (Lla-mas *et al*., 2016; Moreno-Mayar *et al*., 2018; Raghavan *et al*., 2015). Our estimates accounting for TDR are consistent with this scenario and can be considered as additional evidence for this timeframe of migration. Kocher *et al*. (2021) have previously pointed out that a TDR model can recover deeper divergence times for HBV relative to an uncorrelated relaxed clock model, but noted that the standard TDR model (Membrebe *et al*., 2019) provides a poorer fit to the HBV data. Extending a TDR model with branch-specific rates, as we do here using random-effects, is therefore key in providing credibility for the deeper divergence times.

We observed that the TDR is somewhat underestimated relative to estimates for other viruses and that using a prior specification based on these results in better compatibility between divergence times and historical migrations. In this respect, it is important to note that the combined modern and ancient HBV data represent, to our knowledge, the first virus example for which the TDR effect can be estimated from the dated tip structure. Previous analyses of foamy viruses (Aiewsakun and Katzourakis, 2015; Membrebe *et al*., 2019) and lentiviruses (Membrebe *et al*., 2019) involved internal node calibrations over much deeper time scales. Accurately quantifying the TDR effect from dated tips over more limited time scales is likely more challenging, in particular when also confronting other sources of evolutionary variation.

We also considered host-specific rates of evolution by al-lowing a rate effect in the two non-human primate lineages. This revealed a considerably lower non-human primate rate of evolution in the fixed-effects model, but it did not hold up in the mixed-effects model. This may be attributed to an identifiability problem as a lower root-to-tip divergence of these lineages can also be accommodated by negative random-effects on their ancestral branches. Unfortunately, we are currently lacking substantial sampling of non-human primate HBV viruses over a broad time range to accurately estimate the evolutionary rate within the non-human primate lineages specifically. More research is therefore needed to characterize the differences in tempo and mode of HBV evolution between humans and non-human primates.

Our branch-specific implementation of the TDR model presents a challenging inference problem because evolutionary rates are modeled as a function of time, which itself must be estimated. This requires jointly estimating highly correlated node heights and branch-specific rates, a task for which univariate MCMC transition kernels fall short. By employing HMC to estimate both branch rates and node heights within our Bayesian framework, we were able to overcome this limitation. Our approach will likely prove useful to provide evolutionary insights into other pathogens. Ancient genomes have also been recovered for parvoviruses, and similar to HBV, this revealed differences between long-term and short-term evolutionary rates (Mühlemann *et al*., 2018b). For hantaviruses, timescale discrepancies have been revealed by contrasting dated-tip estimates with calibration based on spatial information (Saxenhofer *et al*., 2017).

Not only does our model reconcile a narrative of contradictions, but it also sheds new light on HBV dispersal in the context of ancient human migrations. Although previously associated with European farmers (Kocher *et al*., 2021), the origin of the WENBA lineage had remained elusive. Our estimates of dispersal velocity align HBV spread with Neolithic migrations from Near Eastern centers of early agriculture and position viral evolution as a complementary source of evidence alongside archaeological estimates (Ammerman and Cavalli-Sforza, 1971; Pinhasi *et al*., 2005). Importantly, the association between Hepatitis B virus (HBV) spread and the Neolithic expansion lends independent support to a demic component in the spread of farming across Europe. Although radiocarbon-based reconstructions point to substantial regional heterogeneity, ranging from rapid population replacement to more gradual cultural adoption (Gkiasta *et al*., 2003), the dispersal of a virus necessarily entails the physical movement of infected hosts. Given that long-term HBV persistence is strongly shaped by perinatal and early-life transmission, resulting in quasi-vertical transmission across generations, its historical spread is expected to track sustained population movement rather than transient contact alone. The correspondence between HBV phylogeography and the Neolithic expansion therefore supports scenarios in which migrating farming populations played a significant role (Pinhasi *et al*., 2005), in line with the spatiotemporal spread of so-called Anatolian Neolithic Farmer (ANF) ancestry into Europe, particularly in regions marked by rapid demographic turnover, rather than a purely cultural diffusion of agricultural practices.

The more recent Bronze Age expansions of steppe-derived populations represent one of the largest demographic turnover events in Eurasian prehistory, and we link this process to the modern predominance of HBV genotype D across much of Western and Central Eurasia. The faster rate of HBV dispersal during these expansions broadly accords with genetic estimates of the Yamnaya migration front speed, which suggest a rate of approximately 4.2 km/yr (95% CI: 3.5–5.2), although this estimate varies depending on the ancestry cutoff (Racimo *et al*., 2020). While horse-based mobility has often been proposed as a key driver of steppe expansions (Anthony, 2007; Anthony *et al*., 2026), recent ancient DNA analyses of horses indicate that widespread horseback riding emerged only around 4,150 years BP (Librado *et al*., 2024). This timing is further supported by evidence for a selective sweep associated with traits important for horseback riding between 4,650 and 4,150 years BP (Liu *et al*., 2025). Rather than horseback riding, earlier steppe expansions may have been facilitated by technological innovations such as cattle-drawn wheeled transport (Hosek *et al*., 2024), a scenario that may also be more consistent with the modest increase in dispersal rates we infer here. As we estimated higher dispersal rates associated with more recent migration phases, horses may have contributed more to these dynamics, consistent with the timing of the DOM2 horse expansion—the lineage ancestral to nearly all modern domestic horses—which followed the major Yamnaya expansion phase (Librado *et al*., 2024). We note that a recent reinterpretation of genetic and archaeological evidence suggests an earlier emergence of horseback riding, although involving only a limited fraction of the human subjects and spreading primarily within, rather than between, subregions such as the Carpathian Basin and the Pontic–Caspian and Central Asian steppe (Anthony *et al*., 2026). Future research on horse and cattle domestication and dispersal will clarify whether cattle-drawn wagons, horse-drawn chariots, or both were the primary drivers of the expansion of HBV genotype D documented here.

We demonstrate that the historical overlap of WENBA and genotype D in Central Europe gave rise to a recombinant lineage, namely genotype E. Given that genotype E is now a dominant HBV genotype in sub-Saharan Africa—particularly in West and Central Africa (Ingasia *et al*., 2020)—it would have been reasonable to assume that this lineage emerged on that continent. However, our findings instead illustrate that detailed historical reconstructions are essential for interpreting the present-day geographical distribution of HBV diversity, and that recombination events can contribute important information. The spread of HBV to Europe through both Neolithic and steppe migrations parallels a recent scenario proposed for the origin of Indo-European languages (Heggarty *et al*., 2023). This origin has been the subject of longstanding debate (e.g. Bouckaert *et al*. (2012); Haak *et al*. (2015), with the Anatolian hypothesis linking it to Neolithic farmers and the Steppe hypothesis attributing it to Bronze Age migrations from the Pontic-Caspian steppe. Recent analysis of linguistic divergence has proposed a hybrid hypothesis, with a primary origin in the Fertile Crescent and a secondary homeland on the Pontic-Caspian steppe (Heggarty *et al*., 2023). Our analy-ses indicate that both the Neolithic and steppe migrations also contributed to the spread of HBV to Europe.

Taken together, our results demonstrate that small viral genomes can retain a rich record of past evolutionary and demographic events. When analyzed with models that appropriately capture the complexities of rate variation, ancient HBV sequences become a powerful window into human history and the processes that shaped viral diversity.

## Acknowledgements

We thank Oliver Laeyendecker and Thomas Quinn (Johns Hopkins University) for providing serum samples from patients at the Nioki Hospital (Nioki, DRC). The research leading to these results has received funding from the European Research Council under the European Union’s Horizon 2020 research and innovation programme (grant agreement no. 725422-ReservoirDOCS) and from the European Union’s Horizon 2020 project MOOD (grant agreement no. 874850). PL acknowledges support by the Research Foundation - Flanders (‘Fonds voor Wetenschappelijk Onderzoek - Vlaanderen’, G010326N and G051322N). BV is a postdoctoral researcher of the Fonds de la Recherche Scientifique – FNRS. GB acknowledges support from the Research Foundation - Flanders (Fonds voor Wetenschappelijk Onderzoek -Vlaan-deren,’ G098321N), from the European Union Horizon 2023 RIA project LEAPS (grant agreement no. 101094685) and from the DURABLE EU4Health project 02/2023-01/2027, which is co-funded by the European Union (call EU4H-2021-PJ4) under Grant Agreement No. 101102733. LEK acknowledges support by the Research Foundation - Flanders (‘Fonds voor Wetenschappelijk Onderzoek - Vlaanderen’, 12 × 9222N and G005323N). MAS acknowledges support from National Institutes of Health grants U19 AI135995 and R01 AI153044. We also gratefully acknowledge support from AMD, Inc., with the donation of parallel computing resources used for this research.

