## Supplementary information for "Reconciling fast Hepatitis B evolutionary rates with ancient co-divergence"

### Supporting Information for
### ‘Reconciling fast Hepatitis B evolutionary rates with ancient co-divergence’

Philippe Lemey∗1, Xiang Ji∗2, Bram Vrancken∗1, 3, Magdalini Bletsa∗4, Pratyusa Datta∗5, Liana E. Kafetzopoulou∗1, Jonathon C.O. Mifsud∗1, Guy Baele∗1, Mahmoud Reza Pourkarim∗1, Livia Patrono∗6, 7, Sebastien Calvignac-Spencer∗6, 7, 8, Ludovic Orlando∗9, Paul Bastide∗10, Stephane Guindon∗11, Darren Martin∗12, Marc A. Suchard∗5, 13, 14

∗1Department of Microbiology and Immunology, Rega Institute, KU Leuven – University of Leuven, Leuven, Belgium
∗2Department of Statistics, College of Liberal Arts and Sciences, Iowa State University, Ames, IA, USA
∗3Spatial Epidemiology Lab (SpELL), Université Libre de Bruxelles, Bruxelles, Belgium
∗4Bioinformatics and Applied Genomics Unit, Department of Microbiology, Hellenic Pasteur Institute, Athens, Greece
∗5Department of Biostatistics, Fielding School of Public Health, University of California, Los Angeles, CA, USA
∗6Epidemiology of Highly Pathogenic Microorganisms, Robert Koch Institute, Berlin, Germany
∗7Helmholtz Institute for One Health, Helmholtz-Centre for Infection Research (HZI), Greifswald, Germany
∗8Faculty of Mathematics and Natural Sciences, University of Greifswald, Greifswald, Germany
∗9Centre d’Anthropobiologie et de Génomique de Toulouse (CNRS UMR 5288), Faculté de Santé, Université de Toulouse, Toulouse, France
∗10Université Paris Cité, CNRS, MAP5, F-75006 Paris, France
∗11Équipe Méthodes et Algorithmes pour la Bioinformatique, Laboratoire d’Informatique, de Robotique et de Microélectronique de Montpellier, CNRS—UMR 5506, Montpellier
∗12Faculty of Health Sciences, Institute of Infectious Disease and Molecular Medicine & Department of Integrative Biomedical Sciences, Computational Biology Division, University of Cape Town, Cape Town, South Africa
∗13Department of Biomathematics, David Geffen School of Medicine, University of California, Los Angeles, CA, USA
∗14Department of Human Genetics, David Geffen School of Medicine, University of California, Los Angeles, CA, USA

#### Whole genome sequencing and assembly of specimen HS168

Serum specimens were obtained from patients at the Nioki Hospital (Nioki, DRC) between May 1988 and 1990, as part of an epidemiological study on the impact of Human Immunodeficiency virus 1 (HIV-1) infection in the treatment of patients with African trypanosomiasis [Pepin et al., 1992]. We revisited this collection in order to obtain genomic insights into the microbiota present in those specimens.

Briefly, for specimen HS168, viral RNA was purified with the QIAamp Viral RNA Mini kit (Qiagen) and elution was performed twice using 60 µL and 40 µL of RNase free water, respectively. Reverse transcription and second strand synthesis were performed using random primers and the Superscript IV Reverse Transcriptase enzyme (ThermoFisher Scientific), followed by the NEBNext® Ultra II Non-Directional RNA Second Strand Synthesis Module (New England BioLabs). We prepared sequencing libraries with the NEBNext Ultra II DNA Library Prep Kit for Illumina (New England BioLabs) according to manufacturer's guidelines. Quality was assessed on a BioAnalyzer 2100 using the Agilent High Sensitivity DNA chip (Agilent Technologies) and libraries were quantified using the KAPA Library Quantification kit for Illumina (Roche). Paired-end sequencing was launched by the Nucleomics Core (VIB, Leuven, Belgium) on an Illumina MiSeq platform (PE 300, v3 kit, Illumina).

Following adapter removal using Trimmomatic v0.36 [Bolger et al., 2014], contigs were generated with three different tools: SPAdes v3.12.0 [Bankevich et al., 2012], Ray v2.3.1 [Boisvert et al., 2010] and MEGAHIT v1.2.9 [Li et al., 2015]. All the assembled contigs were analyzed using MegaBLAST [Morgulis et al., 2008] against a virus enriched database. Upon a positive HBV hit, the closest HBV genome was retrieved from GenBank and the trimmed reads were mapped against it using Bowtie2 v2.2.5 [Langmead and Salzberg, 2012]. Coverage and sequencing depth were assessed by calculating the proportion of the mapped reads over the total numbers of reads using SAMtools v1.5 [Li et al., 2009] and BEDtools v2.27.1 [Quinlan and Hall, 2010]. Open reading frames were predicted in Geneious Prime v2019.2.1 [Biomatters, 2019] based on the HBV reference genome (GenBank accession number: NC_003977). The mean sequencing depth of HS168 was 668x and 2 187 positions out of 3 203 had a coverage larger than 20x (68.3%). The HBV genomic sequence generated in this study was deposited in GenBank under accession number MZ005205.

#### The clock model in the partial vectors for one branch

The partial differential equation

$$\frac{d\mathbf{P}(t)}{dt} = \mathbf{P}(t)\mathbf{Q}\mu(t)$$

has the solution involving matrix exponential for one branch of time $(t_1, t_2)$:

$$\log \mathbf{P}(t_2) - \log \mathbf{P}(t_1) = \int_{t_1} *t_2 \mathbf{Q}\mu(t)dt$$

1. For constant $\mu(t) = r$.

$$\mathbf{P}(t_2) = \mathbf{P}(t_1)e * \mathbf{Q}r(t_2 - t_1)$$

that translates to

$$r(t_1, t_2) = r$$

2. For a linear (i.e. time dependent) function, $\mu(t) = \beta t$

$$\mathbf{P}(t_2) = \mathbf{P}(t_1)e * \frac{1}{2}\mathbf{Q}\beta(t_2 * 2 - t_1 * 2)$$
$$= \mathbf{P}(t_1)e * \mathbf{Q}(t_2 - t_1)\beta\frac{t_1 + t_2}{2}$$

which suggests a mid-point for node heights with

$$r(t_1, t_2) = \beta\frac{t_1 + t_2}{2}$$

3. For a linear function with a upper threshold,

$$\mu(t) = \begin{cases} \beta t & t \geq T \\ \beta T & t < T \end{cases}$$

has

$$\mathbf{P}(t_2) = \begin{cases} \mathbf{P}(t_1)e * \mathbf{Q}\beta T(t_2 - t_1) & t_1 < t_2 < T \\ \mathbf{P}(t_1)e * \mathbf{Q}\beta T(T - t_1) + \mathbf{Q}(t_2 - T)\beta\frac{t_2+T}{2} & t_1 < T < t_2 \\ \mathbf{P}(t_1)e * \mathbf{Q}(t_2 - t_1)\beta\frac{t_1+t_2}{2} & T < t_1 < t_2 \end{cases}$$

that translates into

$$\mathbf{r}(t_1, t_2) = \begin{cases} \beta T & t_1 < t_2 < T \\ \beta\frac{\frac{1}{2}(t_2 * 2 - T * 2) + T(T - t_1)}{t_2 - t_1} & t_1 < T < t_2 \\ \beta\frac{t_1+t_2}{2} & T < t_1 < t_2 \end{cases}$$

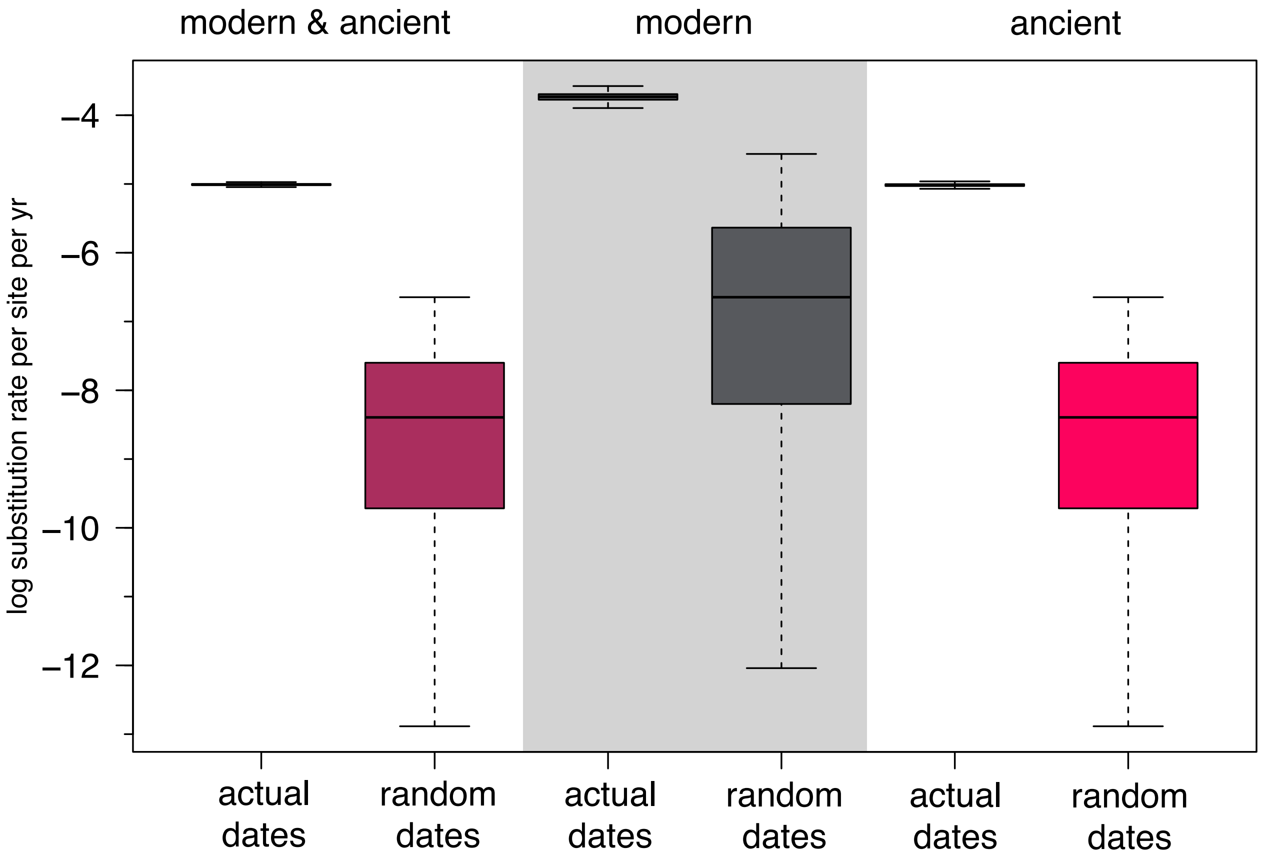


Figure 1: **Date randomisation test to assess temporal signal.** The box plots represent posterior rate distributions under a dated tip model based on the actual dates as well as dates randomized during the inference procedure. The estimates are shown for the complete (modern & ancient) genomes, the modern only genomes and the ancient only genomes.

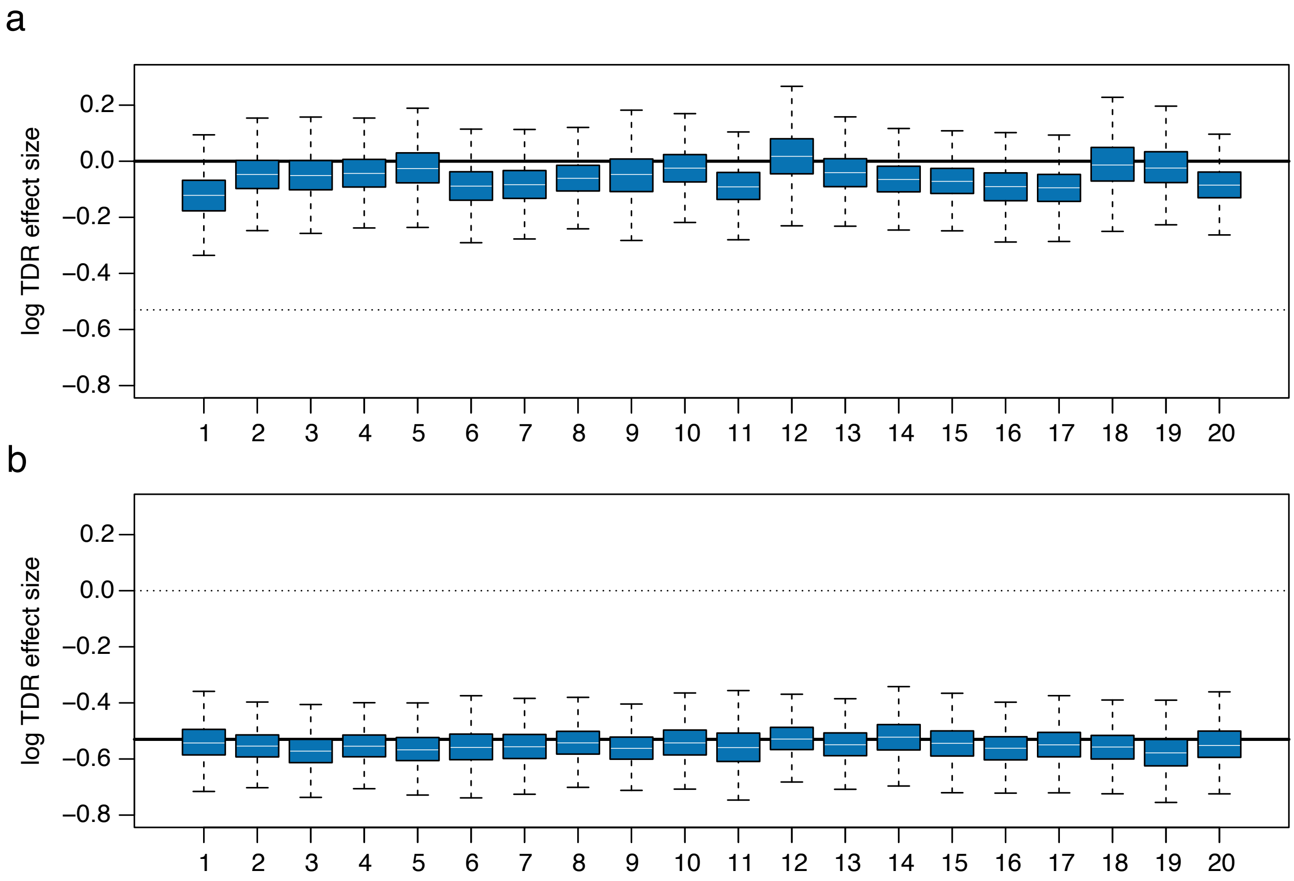


Figure 2: **Time-dependent rate effect estimates for two different simulation scenarios.** Both plots show log time-dependent rate (TDR) effects estimated using a branch-specific TDR model with random-effects fitted to 20 simulation replicates. a) simulations using an uncorrelated relaxed clock model (without TDR effect). b) simulations using an epoch model [Membrebe et al., 2019] with TDR effect (-0.53).

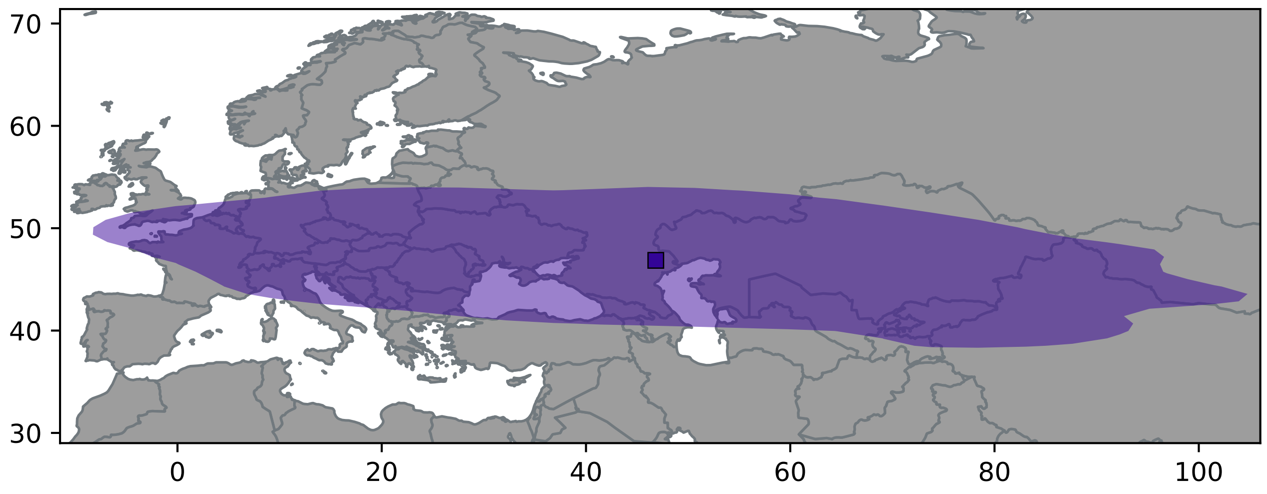


Figure 3: **Spatial projection of the root location in the maximum clade credibility tree for the relaxed random walk analysis of genotype D.** The square represents the mean root locations while the polygon represents the 80% highest density posterior interval.

Table 1: HBV sequences with metadata. For each sequence, we list the accession number if available, the host, the genotype if available, the HBeAg state and the Age. The later can be a fixed year or an age distribution.

| Sequence | Accession | Host | Genotype | HBeAg | Age |
|---|---|---|---|---|---|
| AB194950\|Cameroon\|A\|ep\|19 | AB194950 | human | A | + | 19 |
| AB194951\|Cameroon\|A\|ep\|19 | AB194951 | human | A | + | 19 |
| AB194952\|Cameroon\|A\|ep\|19 | AB194952 | human | A | + | 19 |
| AM184125\|Gabon\|A\|ep\|12 | AM184125 | human | A | + | 12 |
| AM184126\|Gabon\|A\|ep\|12 | AM184126 | human | A | + | 12 |
| AP007263\|Japan\|A\|ep\|12 | AP007263 | human | A | + | 12 |
| AY934763\|Gambia\|A\|ep\|31 | AY934763 | human | A | + | 31 |
| AY934764\|Gambia\|A\|ep\|21 | AY934764 | human | A | + | 21 |
| AY934766\|Somalia\|A\|ep\|18 | AY934766 | human | A | + | 18 |
| AY934767\|Somalia\|A\|ep\|17 | AY934767 | human | A | + | 17 |
| AY934768\|Somalia\|A\|ep\|18 | AY934768 | human | A | + | 18 |
| AY934769\|Somalia\|A\|ep\|18 | AY934769 | human | A | + | 18 |
| AY934770\|Somalia\|A\|ep\|17 | AY934770 | human | A | + | 17 |
| AY934771\|Somalia\|A\|ep\|17 | AY934771 | human | A | + | 17 |
| AY934772\|Uganda\|A\|ep\|12 | AY934772 | human | A | + | 12 |
| AY934773\|Tanzania\|A\|ep\|19 | AY934773 | human | A | + | 19 |
| AY934774\|Philippines\|A\|ep\|17 | AY934774 | human | A | + | 17 |
| DQ020002\|Congo\|A\|ep\|10 | DQ020002 | human | A | + | 10 |
| DQ020003\|UnitedArabEmirates\|A\|ep\|25 | DQ020003 | human | A | + | 25 |
| HS168\|DRC\|A\|ep\|24 | MZ005205 | human | A | + | $\mathcal{U}(23, 25)$ |
| FJ692554\|Nigeria\|A3\|ep\|9 | FJ692554 | human | A3 | + | 9 |
| FJ692555\|Nigeria\|A3\|ep\|9 | FJ692555 | human | A3 | + | 9 |
| FJ692556\|Nigeria\|A3\|ep\|9 | FJ692556 | human | A3 | + | 9 |
| FJ692593\|Haiti\|A3\|ep\|7 | FJ692593 | human | A3 | + | 7 |
| FJ692594\|Haiti\|A3\|ep\|7 | FJ692594 | human | A3 | + | 7 |
| FJ692595\|Haiti\|A3\|ep\|7 | FJ692595 | human | A3 | + | 7 |
| FJ692596\|Haiti\|A3\|ep\|7 | FJ692596 | human | A3 | + | 7 |
| FJ692597\|Haiti\|A3\|ep\|7 | FJ692597 | human | A3 | + | 7 |
| FJ692598\|Haiti\|A3\|ep\|7 | FJ692598 | human | A3 | + | 7 |
| FJ692599\|Haiti\|A3\|ep\|7 | FJ692599 | human | A3 | + | 7 |
| FJ692600\|Haiti\|A3\|ep\|7 | FJ692600 | human | A3 | + | 7 |
| FJ692602\|Haiti\|A3\|ep\|7 | FJ692602 | human | A3 | + | 7 |
| FJ692603\|Haiti\|A3\|ep\|7 | FJ692603 | human | A3 | + | 7 |
| FJ692604\|Haiti\|A3\|ep\|7 | FJ692604 | human | A3 | + | 7 |
| FJ692605\|Haiti\|A3\|ep\|7 | FJ692605 | human | A3 | + | 7 |
| FJ692606\|Haiti\|A3\|ep\|7 | FJ692606 | human | A3 | + | 7 |
| FJ692607\|Haiti\|A3\|ep\|7 | FJ692607 | human | A3 | + | 7 |
| FJ692608\|Haiti\|A3\|ep\|7 | FJ692608 | human | A3 | + | 7 |
| FJ692609\|Haiti\|A3\|en\|7 | FJ692609 | human | A3 | - | 7 |
| FJ692611\|Haiti\|A3\|ep\|7 | FJ692611 | human | A3 | + | 7 |
| FJ692613\|Haiti\|A3\|ep\|7 | FJ692613 | human | A3 | + | 7 |
| AB010289\|Japan\|B1\|ep\|20 | AB010289 | human | B1 | + | 20 |
| AB010290\|Japan\|B1\|en\|20 | AB010290 | human | B1 | - | 20 |
| AB010291\|Japan\|B1\|en\|20 | AB010291 | human | B1 | - | 20 |

*Continued on next page*

| Sequence | Accession | Host | Genotype | HBeAg | Age |
|---|---|---|---|---|---|
| AB010292\|Japan\|B1\|en\|20 | AB010292 | human | B1 | - | 20 |
| AB073838\|Japan\|B1\|en\|12 | AB073838 | human | B1 | - | 12 |
| AB287314\|Alaska\|B5\|ep\|40 | AB287314 | human | B5 | + | 40 |
| AB287315\|Alaska\|B5\|ep\|38 | AB287315 | human | B5 | + | 38 |
| AB287316\|Alaska\|B5\|en\|40 | AB287316 | human | B5 | - | 40 |
| AB287317\|Alaska\|B5\|en\|39 | AB287317 | human | B5 | - | 39 |
| AB287318\|Alaska\|B5\|en\|9 | AB287318 | human | B5 | - | 9 |
| AB287319\|Alaska\|B5\|en\|9 | AB287319 | human | B5 | - | 9 |
| AB287320\|WestGreenland\|B5\|en\|15 | AB287320 | human | B5 | - | 15 |
| AB287321\|WestGreenland\|B5\|en\|15 | AB287321 | human | B5 | - | 15 |
| AB287322\|WestGreenland\|B5\|en\|9 | AB287322 | human | B5 | - | 9 |
| AB287323\|WestGreenland\|B5\|en\|15 | AB287323 | human | B5 | - | 15 |
| AB287324\|WestGreenland\|B5\|en\|9 | AB287324 | human | B5 | - | 9 |
| AB287325\|WestGreenland\|B5\|en\|9 | AB287325 | human | B5 | - | 9 |
| AB287326\|Japan\|B1\|en\|7 | AB287326 | human | B1 | - | 7 |
| AB287327\|Japan\|B1\|en\|7 | AB287327 | human | B1 | - | 7 |
| AB602818\|Japan\|B1\|ep\|8 | AB602818 | human | B1 | + | 8 |
| D23677\|Japan\|B1\|ep\|13 | D23677 | human | B1 | + | 13 |
| D23678\|Japan\|B1\|en\|13 | D23678 | human | B1 | - | 13 |
| D23679\|Japan\|B1\|en\|13 | D23679 | human | B1 | - | 13 |
| DQ463795\|WestNunavut\|B5\|en\|9 | DQ463795 | human | B5 | - | 9 |
| DQ463796\|WestNunavut\|B5\|en\|9 | DQ463796 | human | B5 | - | 9 |
| DQ463799\|WestNunavut\|B5\|en\|9 | DQ463799 | human | B5 | - | 9 |
| DQ463802\|WestNunavut\|B5\|en\|9 | DQ463802 | human | B5 | - | 9 |
| FJ386584\|China\|B2\|ep\|5 | FJ386584 | human | B2 | + | 5 |
| FJ386600\|China\|B2\|ep\|5 | FJ386600 | human | B2 | + | 5 |
| FJ386636\|China\|B2\|ep\|5 | FJ386636 | human | B2 | + | 5 |
| GQ924653\|Malaysia\|B2\|ep\|6 | GQ924653 | human | B2 | + | 6 |
| JN792894\|WestNunavut\|B5\|en\|4 | JN792894 | human | B5 | - | 4 |
| JN792896\|WestNunavut\|B5\|en\|4 | JN792896 | human | B5 | - | 4 |
| JN792897\|WestNunavut\|B5\|en\|4 | JN792897 | human | B5 | - | 4 |
| JN792900\|WestNunavut\|B5\|en\|4 | JN792900 | human | B5 | - | 4 |
| JN792901\|WestNunavut\|B5\|en\|4 | JN792901 | human | B5 | - | 4 |
| KP659219\|Alaska\|B5\|en\|7 | KP659219 | human | B5 | - | 7 |
| KP659220\|Alaska\|B5\|en\|8 | KP659220 | human | B5 | - | 8 |
| KP659221\|Alaska\|B5\|en\|8 | KP659221 | human | B5 | - | 8 |
| KP659223\|Alaska\|B5\|en\|10 | KP659223 | human | B5 | - | 10 |
| KP659224\|Alaska\|B5\|en\|5 | KP659224 | human | B5 | - | 5 |
| KP659234\|WestNunavut\|B5\|ep\|30 | KP659234 | human | B5 | + | 30 |
| KP659235\|WestNunavut\|B5\|ep\|30 | KP659235 | human | B5 | + | 30 |
| KP659237\|EastNunavut\|B5\|ep\|30 | KP659237 | human | B5 | + | 30 |
| KP659245\|EastNunavut\|B5\|en\|1 | KP659245 | human | B5 | - | 1 |
| KP659246\|EastNunavut\|B5\|en\|30 | KP659246 | human | B5 | - | 30 |
| KP659247\|EastNunavut\|B5\|ep\|30 | KP659247 | human | B5 | + | 30 |
| KP659248\|EastNunavut\|B5\|en\|0 | KP659248 | human | B5 | - | 0 |
| KP659249\|EastNunavut\|B5\|en\|0 | KP659249 | human | B5 | - | 0 |
| KP659250\|EastNunavut\|B5\|en\|0 | KP659250 | human | B5 | - | 0 |
| KP659251\|EastNunavut\|B5\|en\|0 | KP659251 | human | B5 | - | 0 |

*Continued on next page*

| Sequence | Accession | Host | Genotype | HBeAg | Age |
|---|---|---|---|---|---|
| KP659252\|EastNunavut\|B5\|en\|0 | KP659252 | human | B5 | - | 0 |
| KP659253\|EastNunavut\|B5\|en\|0 | KP659253 | human | B5 | - | 0 |
| KP659254\|EastNunavut\|B5\|en\|0 | KP659254 | human | B5 | - | 0 |
| KP659255\|WestNunavut\|B5\|en\|0 | KP659255 | human | B5 | - | 0 |
| AB014360\|01D01HCC\|C\|ep\|18 | AB014360 | human | C | + | 18 |
| AB014362\|03D03HCC\|C\|ep\|17 | AB014362 | human | C | + | 17 |
| AB014363\|04D04HCC\|C\|ep\|17 | AB014363 | human | C | + | 17 |
| AB014367\|08D08HCC\|C\|ep\|24 | AB014367 | human | C | + | 24 |
| AB014380\|21Y03HCC\|C\|ep\|25 | AB014380 | human | C | + | 25 |
| AB014391\|32Y14HCC\|C\|ep\|23 | AB014391 | human | C | + | 23 |
| AB014394\|35Y17HCC\|C\|ep\|17 | AB014394 | human | C | + | 17 |
| AB113879\|Japan\|C\|ep\|15 | AB113879 | human | C | + | 15 |
| AF223954\|EastAsia\|C\|ep\|25 | AF223954 | human | C | + | 25 |
| AF223955\|EastAsia\|C\|ep\|24 | AF223955 | human | C | + | 24 |
| AF223957\|EastAsia\|C\|ep\|27 | AF223957 | human | C | + | 27 |
| AF223958\|EastAisa\|C\|ep\|24 | AF223958 | human | C | + | 24 |
| AF223956\|EastAsia\|C\|ep\|26 | AF223956 | human | C | + | 26 |
| AF223960\|EastAsia\|C\|ep\|28 | AF223960 | human | C | + | 28 |
| AF223961\|EastAsia\|C\|ep\|22 | AF223961 | human | C | + | 22 |
| AY247030\|Korea\|C\|ep\|15 | AY247030 | human | C | + | 15 |
| AY247031\|Korea\|C\|ep\|16 | AY247031 | human | C | + | 16 |
| AY167090\|Taiwan\|C\|ep\|15 | AY167090 | human | C | + | 15 |
| AY167091\|Tiawan\|C\|ep\|17 | AY167091 | human | C | + | 17 |
| AY167092\|Taiwan\|C\|epi\|16 | AY167092 | human | C | +i | 16 |
| HQ700498\|Fiji\|C\|ep\|8 | HQ700498 | human | C | + | 8 |
| HQ700495\|Fiji\|C\|ep\|9 | HQ700495 | human | C | + | 9 |
| HQ700502\|TONGA\|C\|ep\|22 | HQ700502 | human | C | + | 22 |
| HQ700504\|TONGA\|C\|ep\|21 | HQ700504 | human | C | + | 21 |
| HQ700505\|VANUATA\|C\|ep\|9 | HQ700505 | human | C | + | 9 |
| HQ700506\|N_CALEDONIA\|C\|ep\|10 | HQ700506 | human | C | + | 10 |
| HQ700507\|N_CALEDONIA\|C\|ep\|12 | HQ700507 | human | C | + | 12 |
| HQ700508\|N_CALEDONIA\|C\|ep\|10 | HQ700508 | human | C | + | 10 |
| HQ700509\|N_CALEDONIA\|C\|ep\|10 | HQ700509 | human | C | + | 10 |
| HQ700523\|PNG\|C\|ep\|9 | HQ700523 | human | C | + | 9 |
| HQ700527\|PNG\|C\|ep\|9 | HQ700527 | human | C | + | 9 |
| HQ700519\|PNG\|C\|ep\|10 | HQ700519 | human | C | + | 10 |
| HQ700520\|PNG\|C\|ep\|9 | HQ700520 | human | C | + | 9 |
| HQ700521\|PNG\|C\|ep\|9 | HQ700521 | human | C | + | 9 |
| HQ700522\|PNG\|C\|ep\|9 | HQ700522 | human | C | + | 9 |
| HQ700516\|PNG\|C\|ep\|8 | HQ700516 | human | C | + | 8 |
| HQ700518\|PNG\|C\|ep\|8 | HQ700518 | human | C | + | 8 |
| HQ700530\|PNG\|C\|ep\|19 | HQ700530 | human | C | + | 19 |
| HQ700531\|PNG\|C\|ep\|19 | HQ700531 | human | C | + | 19 |
| HQ700532\|PNG\|C\|ep\|19 | HQ700532 | human | C | + | 19 |
| HQ700456\|NZL\|C\|ep\|29 | HQ700456 | human | C | + | 29 |
| HQ700485\|NZL\|C\|ep\|29 | HQ700485 | human | C | + | 29 |
| HQ700461\|NZL\|C\|ep\|29 | HQ700461 | human | C | + | 29 |
| HQ700526\|PNG\|C\|ep\|8 | HQ700526 | human | C | + | 8 |

*Continued on next page*

| Sequence | Accession | Host | Genotype | HBeAg | Age |
|---|---|---|---|---|---|
| HQ700517\|PNG\|C\|ep\|8 | HQ700517 | human | C | + | 8 |
| AB014365\|Japan\|C\|en\|25 | AB014365 | human | C | - | 25 |
| AB014379\|Japan\|C\|en\|25 | AB014379 | human | C | - | 25 |
| AB014381\|Japan\|C\|en\|25 | AB014381 | human | C | - | 25 |
| AB014382\|Japan\|C\|en\|23 | AB014382 | human | C | - | 23 |
| AB014383\|Japan\|C\|en\|23 | AB014383 | human | C | - | 23 |
| AB014384\|Japan\|C\|en\|23 | AB014384 | human | C | - | 23 |
| AB014385\|Japan\|C\|en\|23 | AB014385 | human | C | - | 23 |
| AB014392\|Japan\|C\|en\|23 | AB014392 | human | C | - | 23 |
| AB014393\|Japan\|C\|en\|18 | AB014393 | human | C | - | 18 |
| AB014396\|Japan\|C\|en\|17 | AB014396 | human | C | - | 17 |
| AB049609\|Japan\|C\|en\|17 | AB049609 | human | C | - | 17 |
| AB049610\|Japan\|C\|en\|17 | AB049610 | human | C | - | 17 |
| D23682\|Japan\|C\|en\|29 | D23682 | human | C | - | 29 |
| D23684\|Japan\|C\|en\|25 | D23684 | human | C | - | 25 |
| AB113878\|Japan\|C\|en\|13 | AB113878 | human | C | - | 13 |
| AF330110\|EastAsia\|C\|en\|16 | AF330110 | human | C | - | 16 |
| HQ700462\|NZL\|C\|en\|17 | HQ700462 | human | C | - | 17 |
| HQ700490\|NZL\|C\|en\|20 | HQ700490 | human | C | - | 20 |
| AF043593\|Germany\|D\|ep\|24 | AF043593 | human | D | + | 24 |
| AY741797\|Iran\|D\|ep\|10 | AY741797 | human | D | + | 10 |
| AB090268\|Japan\|D\|ep\|16 | AB090268 | human | D | + | 16 |
| AB090269\|Japan\|D\|ep\|21 | AB090269 | human | D | + | 21 |
| AB078033\|Japan\|D\|ep\|15 | AB078033 | human | D | + | 15 |
| AB090270\|Japan\|D\|ep\|14 | AB090270 | human | D | + | 14 |
| AB109475\|Japan\|D\|ep\|12 | AB109475 | human | D | + | 12 |
| AB109476\|Japan\|D\|ep\|16 | AB109476 | human | D | + | 16 |
| AB120308\|Japan\|D\|ep\|31 | AB120308 | human | D | + | 31 |
| AF121240\|NA\|D\|ep\|19 | AF121240 | human | D | + | 19 |
| HQ700497\|FIJI\|D\|ep\|9 | HQ700497 | human | D | + | 9 |
| HQ700503\|TONGA\|D\|ep\|21 | HQ700503 | human | D | + | 21 |
| HQ700501\|SAMOA\|D\|ep\|22 | HQ700501 | human | D | + | 22 |
| HQ700511\|N_CALEDONIA\|D\|ep\|12 | HQ700511 | human | D | + | 12 |
| HQ700512\|N_CALEDONIA\|D\|ep\|12 | HQ700512 | human | D | + | 12 |
| HQ700513\|N_CALEDONIA\|D\|ep\|12 | HQ700513 | human | D | + | 12 |
| HQ700514\|N_CALEDONIA\|D\|ep\|12 | HQ700514 | human | D | + | 12 |
| HQ700524\|PNG\|D\|ep\|9 | HQ700524 | human | D | + | 9 |
| HQ700525\|PNG\|D\|ep\|9 | HQ700525 | human | D | + | 9 |
| HQ700541\|Kiribati\|D\|ep\|8 | HQ700541 | human | D | + | 8 |
| HQ700538\|Kiribati\|D\|ep\|9 | HQ700538 | human | D | + | 9 |
| HQ700464\|NZL\|D\|ep\|29 | HQ700464 | human | D | + | 29 |
| HQ700442\|NZL\|D\|ep\|14 | HQ700442 | human | D | + | 14 |
| HQ700466\|NZL\|D\|ep\|29 | HQ700466 | human | D | + | 29 |
| HQ700470\|NZL\|D\|ep\|14 | HQ700470 | human | D | + | 14 |
| HQ700474\|NZL\|D\|ep\|14 | HQ700474 | human | D | + | 14 |
| HQ700484\|NZL\|D\|ep\|29 | HQ700484 | human | D | + | 29 |
| HQ700459\|NZL\|D\|ep\|20 | HQ700459 | human | D | + | 20 |
| HQ700478\|NZL\|D\|en\|19 | HQ700478 | human | D | - | 19 |

*Continued on next page*

| Sequence | Accession | Host | Genotype | HBeAg | Age |
|---|---|---|---|---|---|
| HQ700481\|NZL\|D\|en\|14 | HQ700481 | human | D | - | 14 |
| HQ700472\|NZL\|D\|en\|16 | HQ700472 | human | D | - | 16 |
| HQ700458\|NZL\|D\|en\|16 | HQ700458 | human | D | - | 16 |
| HQ700446\|NZL\|D\|ep\|14 | HQ700446 | human | D | + | 14 |
| HQ700448\|NZL\|D\|ep\|14 | HQ700448 | human | D | + | 14 |
| HQ700455\|NZL\|D\|ep\|14 | HQ700455 | human | D | + | 14 |
| AY741794\|Iran\|D\|en\|10 | AY741794 | human | D | - | 10 |
| AY741795\|Iran\|D\|en\|9 | AY741795 | human | D | - | 9 |
| AY741798\|Iran\|D\|en\|9 | AY741798 | human | D | - | 9 |
| AY741796\|Iran\|D\|en\|9 | AY741796 | human | D | - | 9 |
| AB109478\|Japan\|D\|en\|15 | AB109478 | human | D | - | 15 |
| AB109479\|Japan\|D\|en\|16 | AB109479 | human | D | - | 16 |
| AB119251\|Japan\|D\|en\|11 | AB119251 | human | D | - | 11 |
| AB119252\|Japan\|D\|en\|11 | AB119252 | human | D | - | 11 |
| AB119254\|Japan\|D\|en\|10 | AB119254 | human | D | - | 10 |
| AF121239\|Japan\|D\|en\|20 | AF121239 | human | D | - | 20 |
| HQ700540\|Kiribati\|D\|en\|9 | HQ700540 | human | D | - | 9 |
| HQ700533\|Kiribati\|D\|en\|9 | HQ700533 | human | D | - | 9 |
| HQ700534\|Kiribati\|D\|en\|9 | HQ700534 | human | D | - | 9 |
| HQ700535\|Kiribati\|D\|en\|9 | HQ700535 | human | D | - | 9 |
| HQ700537\|Kiribati\|D\|en\|9 | HQ700537 | human | D | - | 9 |
| HQ700536\|Kiribati\|D\|en\|9 | HQ700536 | human | D | - | 9 |
| HQ700510\|N_CALEDONIA\|D\|en\|10 | HQ700510 | human | D | - | 10 |
| HQ700492\|Fiji\|D\|en\|9 | HQ700492 | human | D | - | 9 |
| HQ700493\|Fiji\|D\|en\|9 | HQ700493 | human | D | - | 9 |
| HQ700494\|Fiji\|D\|en\|9 | HQ700494 | human | D | - | 9 |
| HQ700500\|Samoa\|D\|en\|20 | HQ700500 | human | D | - | 20 |
| AB194947\|Cameroon\|E\|ep\|19 | AB194947 | human | E | + | 19 |
| AB205129\|Ghana\|E\|ep\|13 | AB205129 | human | E | + | 13 |
| AB205188\|Ghana\|E\|epi\|13 | AB205188 | human | E | +i | 13 |
| AB205189\|Ghana\|E\|ep\|13 | AB205189 | human | E | + | 13 |
| AB205190\|Ghana\|E\|ep\|13 | AB205190 | human | E | + | 13 |
| AB205191\|Ghana\|E\|ep\|13 | AB205191 | human | E | + | 13 |
| AB205192\|Ghana\|E\|ep\|13 | AB205192 | human | E | + | 13 |
| AP007262\|Zambia\|E\|ep\|12 | AP007262 | human | E | + | 12 |
| DQ060822\|Angola\|E\|ep\|20 | DQ060822 | human | E | + | 20 |
| DQ060823\|Angola\|E\|ep\|20 | DQ060823 | human | E | + | 20 |
| AB036905\|Venezuela\|F\|ep\|23 | AB036905 | human | F | + | 23 |
| AB036910\|Venezuela\|F\|ep\|23 | AB036910 | human | F | + | 23 |
| AB036911\|Venezuela\|F\|ep\|23 | AB036911 | human | F | + | 23 |
| AB036912\|Venezuela\|F\|ep\|23 | AB036912 | human | F | + | 23 |
| AB036914\|Venezuela\|F\|ep\|23 | AB036914 | human | F | + | 23 |
| AB036915\|Venezuela\|F\|ep\|23 | AB036915 | human | F | + | 23 |
| AB036916\|Venezuela\|F\|ep\|23 | AB036916 | human | F | + | 23 |
| AB036919\|Venezuela\|F\|ep\|23 | AB036919 | human | F | + | 23 |
| AB036920\|Venezuela\|F\|ep\|22 | AB036920 | human | F | + | 22 |
| AB116549\|Panama\|F\|ep\|13 | AB116549 | human | F | + | 13 |
| AB116550\|Panama\|F\|ep\|13 | AB116550 | human | F | + | 13 |

*Continued on next page*

| Sequence | Accession | Host | Genotype | HBeAg | Age |
|---|---|---|---|---|---|
| X69798\|Brazil\|F\|ep\|32 | X69798 | human | F | + | 32 |
| AF223962\|Argentina\|F\|ep\|17 | AF223962 | human | F | + | 17 |
| AF223963\|Argentina\|F\|ep\|17 | AF223963 | human | F | + | 17 |
| AF223964\|Argentina\|F\|ep\|17 | AF223964 | human | F | + | 17 |
| AF223965\|Argentina\|F\|ep\|17 | AF223965 | human | F | + | 17 |
| AY179735\|Argintina\|F\|ep\|16 | AY179735 | human | F | + | 16 |
| AY311369\|NA\|F\|ep\|14 | AY311369 | human | F | + | 14 |
| AY311370\|NA\|F\|ep\|14 | AY311370 | human | F | + | 14 |
| AB064316\|USA\|F\|en\|17 | AB064316 | human | F | - | 17 |
| AB166850\|Bolivia\|F\|en\|14 | AB166850 | human | F | - | 14 |
| AB059660\|USA\|H\|ep\|12 | AB059660 | human | H | + | 12 |
| AB375163\|MEX\|H\|ep\|5 | AB375163 | human | H | + | 5 |
| AY090454\|NIC\|H\|ep\|16 | AY090454 | human | H | + | 16 |
| AY090457\|NIC\|H\|ep\|36 | AY090457 | human | H | + | 36 |
| AB486012\|Japan\|J\|en\|7 | AB486012 | human | J | - | 7 |
| AB056513\|USA\|G\|en\|11 | AB056513 | human | G | - | $Exp(\lambda = 2)$ |
| AB064312\|USA\|G\|en\|11 | AB064312 | human | G | - | $Exp(\lambda = 2)$ |
| AF405706\|Germany\|G\|en\|11 | AF405706 | human | G | - | $Exp(\lambda = 2)$ |
| HSJN194\|Mexico\|ep\|465 | MT108214 | human | | + | $\mathcal{N}(\mu = 465, \sigma = 39)$ |
| DA195\|Hungary\|ep\|2641 | LT992441.1 | human | | + | $\mathcal{N}(\mu = 2641, \sigma = 44)$ |
| DA119\|Slovakia\|ep\|1563 | LT992440.1 | human | | + | $\mathcal{U}(1463, 1663)$ |
| RISE386\|Russian_Federation\|ep\|4184 | LT992448 | human | | + | $\mathcal{N}(\mu = 4184, \sigma = 47)$ |
| RISE387\|Russian_Federation\|en\|4278 | LT992447 | human | | - | $\mathcal{N}(\mu = 4278, \sigma = 46)$ |
| JN315779\|South_Korea\|C2\|ep\|333 | JN315779 | human | C2 | + | $\mathcal{N}(\mu = 333, \sigma = 36)$ |
| RISE254\|Hungary\|ep\|4005 | LT992459 | human | | + | $\mathcal{N}(\mu = 4005, \sigma = 43)$ |
| RISE563\|Germany\|ep\|4484 | LT992443 | human | | + | $\mathcal{N}(\mu = 4484, \sigma = 48)$ |
| RISE154\|Poland\|ep\|3847 | LT992455 | human | | + | $\mathcal{N}(\mu = 3847, \sigma = 38)$ |
| DA29\|Kazakhstan\|ep\|818 | LT992438 | human | | + | $\mathcal{N}(\mu = 818, \sigma = 32)$ |
| DA222\|Kazakhstan\|ep\|1163 | LT992454 | human | | + | $\mathcal{U}(1063, 1263)$ |
| DA27\|Kazakhstan\|ep\|1606 | LT992439 | human | | + | $\mathcal{N}(\mu = 1606, \sigma = 40)$ |
| DA51\|Kyrgyzstan\|ep\|2293 | LT992444 | human | | + | $\mathcal{N}(\mu = 2293, \sigma = 45)$ |
| DA45\|Mongolia\|ep\|2116 | LT992442 | human | | + | $\mathcal{N}(\mu = 2116, \sigma = 36)$ |
| MG585269\|Italy\|D3\|ep\|444 | MG585269 | human | D3 | + | $\mathcal{N}(\mu = 444, \sigma = 31)$ |
| Sorsum\|Germany\|en\|5233 | | human | | - | $\mathcal{N}(\mu = 5233, \sigma = 58)$ |
| Karsdorf\|Germany\|en\|7089 | | human | | - | $\mathcal{N}(\mu = 7090, \sigma = 62)$ |
| Petersberg\|Germany\|ep\|931 | | human | | + | $\mathcal{N}(\mu = 932, \sigma = 31)$ |
| SJN001\|Mexico\|A3\|ep\|473 | | human | A3 | + | $\mathcal{N}(\mu = 473, \sigma = 44)$ |
| Abusir1543\|Egypt\|ep\|1924 | | human | | + | $\mathcal{N}(\mu = 1924, \sigma = 18)$ |
| AKB003\|Kazakhstan\|ep\|2649 | | human | | + | $\mathcal{N}(\mu = 2649, \sigma = 67)$ |
| BRE008\|Kazakhstan\|ep\|1701 | | human | | + | $\mathcal{N}(\mu = 1698, \sigma = 32)$ |
| BRE026\|Kazakhstan\|ep\|2363 | | human | | + | $\mathcal{U}(2313, 2413)$ |
| BRE028\|Kazakhstan\|ep\|2363 | | human | | + | $\mathcal{U}(2313, 2413)$ |
| CUN002\|Peru\|ep\|9085 | | human | | + | $\mathcal{N}(\mu = 9086, \sigma = 122)$ |
| I0157\|Great_Britain\|ep\|1295 | | human | | + | $\mathcal{N}(\mu = 1295, \sigma = 24)$ |
| I0161\|Great_Britain\|ep\|1228 | | human | | + | $\mathcal{N}(\mu = 1228, \sigma = 45)$ |
| I0216\|Italy\|ep\|2663 | | human | | + | $\mathcal{U}(2513, 2813)$ |
| I0217\|Italy\|ep\|2663 | | human | | + | $\mathcal{U}(2513, 2813)$ |

*Continued on next page*

| Sequence | Accession | Host | Genotype | HBeAg | Age |
|---|---|---|---|---|---|
| I1344\|Russian_Federation\|ep\|713 | | human | | + | $\mathcal{U}(613, 813)$ |
| KIL044\|Ireland\|en\|838 | | human | | - | $\mathcal{U}(763, 913)$ |
| KRA001\|Germany\|ep\|797 | | human | | + | $\mathcal{N}(\mu = 798, \sigma = 20)$ |
| KRA010\|Germany\|ep\|683 | | human | | + | $\mathcal{N}(\mu = 684, \sigma = 26)$ |
| MAY017\|Russian_Federation\|ep\|663 | | human | | + | $\mathcal{U}(513, 813)$ |
| MLR005\|Russian_Federation\|ep\|713 | | human | | + | $\mathcal{U}(613, 813)$ |
| OAI017\|Great_Britain\|ep\|1513 | | human | | + | $\mathcal{U}(1413, 1613)$ |
| ORE002\|Russian_Federation\|ep\|2585 | | human | | + | $\mathcal{N}(\mu = 2585, \sigma = 81)$ |
| RMI002\|Germany\|ep\|613 | | human | | + | $\mathcal{U}(513, 713)$ |
| SED009\|Great_Britain\|ep\|1263 | | human | | + | $\mathcal{U}(1213, 1313)$ |
| SHK001\|Russian_Federation\|ep\|811 | | human | | + | $\mathcal{N}(\mu = 811, \sigma = 18)$ |
| STR144\|Belgium\|ep\|469 | | human | | + | $\mathcal{N}(\mu = 469, \sigma = 42)$ |
| ZWE008\|Switzerland\|ep\|680 | | human | | + | $\mathcal{N}(\mu = 680, \sigma = 31)$ |
| BNL005\|Czech_Republic\|ep\|4135 | | human | | + | $\mathcal{N}(\mu = 4136, \sigma = 39)$ |
| BON020\|Turkey\|ep\|10095 | | human | | + | $\mathcal{N}(\mu = 10096, \sigma = 67)$ |
| CLL005\|Spain\|ep\|4913 | | human | | + | $\mathcal{U}(4313, 5513)$ |
| DER027\|Germany\|ep\|4013 | | human | | + | $\mathcal{U}(3813, 4213)$ |
| GRG036\|France\|ep\|6763 | | human | | + | $\mathcal{U}(6513, 7013)$ |
| I0061\|Russian_Federation\|ep\|8313 | | human | | + | $\mathcal{U}(7913, 8713)$ |
| I0104\|Germany\|ep\|4435 | | human | | + | $\mathcal{N}(\mu = 4436, \sigma = 63)$ |
| KLE031\|Germany\|ep\|3896 | | human | | + | $\mathcal{N}(\mu = 3896, \sigma = 56)$ |
| Loschbour\|Luxembourg\|ep\|8113 | | human | | + | $\mathcal{N}(\mu = 8114, \sigma = 53)$ |
| MIB003\|Czech_Republic\|ep\|3867 | | human | | + | $\mathcal{N}(\mu = 3868, \sigma = 37)$ |
| MIB040\|Czech_Republic\|ep\|3877 | | human | | + | $\mathcal{N}(\mu = 3877, \sigma = 41)$ |
| MIB041\|Czech_Republic\|ep\|3837 | | human | | + | $\mathcal{N}(\mu = 3838, \sigma = 34)$ |
| MIS002\|Czech_Republic\|ep\|3960 | | human | | + | $\mathcal{N}(\mu = 3960, \sigma = 29)$ |
| MKL025\|Czech_Republic\|ep\|3843 | | human | | + | $\mathcal{N}(\mu = 3843, \sigma = 33)$ |
| MN2003\|Russian_Federation\|ep\|10598 | | human | | + | $\mathcal{N}(\mu = 10598, \sigma = 50)$ |
| MOT001\|Sweden\|ep\|7807 | | human | | + | $\mathcal{N}(\mu = 7808, \sigma = 74)$ |
| MPR001\|Belgium\|ep\|10533 | | human | | + | $\mathcal{N}(\mu = 10534, \sigma = 110)$ |
| OOH025\|Germany\|en\|3646 | | human | | - | $\mathcal{N}(\mu = 3646, \sigma = 46)$ |
| PLZ001\|Spain\|ep\|3638 | | human | | + | $\mathcal{U}(3513, 3763)$ |
| SEC006\|Italy_Sardinia\|ep\|4393 | | human | | + | $\mathcal{N}(\mu = 4393, \sigma = 39)$ |
| SUA002\|Italy_Sardinia\|ep\|4149 | | human | | + | $\mathcal{N}(\mu = 4150, \sigma = 49)$ |
| SUA007\|Italy_Sardinia\|ep\|3969 | | human | | + | $\mathcal{N}(\mu = 3970, \sigma = 30)$ |
| TRI011\|Italy\|ep\|3363 | | human | | + | $\mathcal{U}(3313, 3413)$ |
| UOO033\|Russian_Federation\|ep\|8113 | | human | | + | $\mathcal{U}(8013, 8213)$ |
| UZZ075\|Italy_Sicily\|ep\|7287 | | human | | + | $\mathcal{N}(\mu = 7288, \sigma = 24)$ |
| UZZ081\|Italy_Sicily\|ep\|8649 | | human | | + | $\mathcal{N}(\mu = 8650, \sigma = 20)$ |
| WEH008\|Germany\|ep\|3941 | | human | | + | $\mathcal{N}(\mu = 3942, \sigma = 20)$ |
| WEH009\|Germany\|ep\|3969 | | human | | + | $\mathcal{N}(\mu = 3969, \sigma = 29)$ |
| WSN012\|Germany\|ep\|4198 | | human | | + | $\mathcal{N}(\mu = 4198, \sigma = 78)$ |
| I0461\|Spain\|ep\|4333 | | human | | + | $\mathcal{N}(\mu = 4333, \sigma = 58)$ |
| KVK001\|Russian_Federation\|ep\|10275 | | human | | + | $\mathcal{N}(\mu = 10275, \sigma = 17)$ |
| ELT006\|Spain\|en\|5863 | | human | | - | $\mathcal{N}(\mu = 5863, \sigma = 43)$ |

*Continued on next page*

| Sequence | Accession | Host | Genotype | HBeAg | Age |
|---|---|---|---|---|---|
| HAL05\|Germany\|en\|7105 | | human | | - | $\mathcal{N}(\mu = 7105, \sigma = 56)$ |
| ISB002\|Italy_Sardinia\|en\|3688 | | human | | - | $\mathcal{N}(\mu = 3688, \sigma = 31)$ |
| SID005\|Italy_Sardinia\|en\|6157 | | human | | - | $\mathcal{N}(\mu = 6157, \sigma = 48)$ |
| UZZ099\|Italy_Sicily\|en\|6011 | | human | | - | $\mathcal{N}(\mu = 6011, \sigma = 21)$ |
| VTZ011\|France\|en\|6637 | | human | | - | $\mathcal{N}(\mu = 6637, \sigma = 39)$ |
| I0784\|Bulgaria\|en\|6313 | | human | | - | $\mathcal{U}(6213, 6413)$ |
| ARM001\|Armenia\|ep\|5511 | | human | | + | $\mathcal{N}(\mu = 5511, \sigma = 63)$ |
| ARM0023\|Armenia\|ep\|5195 | | human | | + | $\mathcal{N}(\mu = 5195, \sigma = 75)$ |
| CHA003\|Spain\|ep\|7149 | | human | | + | $\mathcal{N}(\mu = 7149, \sigma = 74)$ |
| ELT005\|Spain\|ep\|7227 | | human | | + | $\mathcal{N}(\mu = 7228, \sigma = 42)$ |
| GGN005\|France\|ep\|5715 | | human | | + | $\mathcal{N}(\mu = 5715, \sigma = 32)$ |
| GRG001\|France\|ep\|7032 | | human | | + | $\mathcal{N}(\mu = 7032, \sigma = 86)$ |
| HAL15\|Germany\|ep\|7055 | | human | | + | $\mathcal{N}(\mu = 7055, \sigma = 68)$ |
| HOP004\|Czech_Republic\|ep\|4310 | | human | | + | $\mathcal{N}(\mu = 4310, \sigma = 63)$ |
| HSL001\|Sweden\|ep\|4365 | | human | | + | $\mathcal{N}(\mu = 4365, \sigma = 111)$ |
| I0100\|Germany\|ep\|7052 | | human | | + | $\mathcal{N}(\mu = 7052, \sigma = 72)$ |
| I0411\|Spain\|ep\|7192 | | human | | + | $\mathcal{N}(\mu = 7192, \sigma = 54)$ |
| I0551\|Germany\|ep\|5213 | | human | | + | $\mathcal{U}(5013, 5413)$ |
| I0795\|Germany\|ep\|7137 | | human | | + | $\mathcal{N}(\mu = 7137, \sigma = 43)$ |
| I0798\|Germany\|ep\|5213 | | human | | + | $\mathcal{U}(5013, 5413)$ |
| I2005\|Germany\|ep\|7187 | | human | | + | $\mathcal{N}(\mu = 7187, \sigma = 57)$ |
| I2020\|Germany\|ep\|7163 | | human | | + | $\mathcal{U}(6813, 7513)$ |
| I2031\|Germany\|ep\|7163 | | human | | + | $\mathcal{U}(6813, 7513)$ |
| IKI009\|Turkey\|ep\|5255 | | human | | + | $\mathcal{N}(\mu = 5256, \sigma = 59)$ |
| JAZ001\|Russian_Federation\|ep\|7307 | | human | | + | $\mathcal{N}(\mu = 7308, \sigma = 30)$ |
| MUR007\|Russian_Federation\|ep\|6472 | | human | | + | $\mathcal{N}(\mu = 6472, \sigma = 45)$ |
| PDA001\|Czech_Republic\|ep\|4069 | | human | | + | $\mathcal{N}(\mu = 4069, \sigma = 38)$ |
| PEN003\|France\|ep\|7419 | | human | | + | $\mathcal{N}(\mu = 7419, \sigma = 36)$ |
| UZZ061\|Italy_Sicily\|ep\|6761 | | human | | + | $\mathcal{N}(\mu = 6761, \sigma = 17)$ |
| VLI016\|Czech_Republic\|ep\|4401 | | human | | + | $\mathcal{N}(\mu = 4401, \sigma = 40)$ |
| VTZ008\|France\|ep\|6413 | | human | | + | $\mathcal{U}(6213, 6613)$ |
| YUN048\|Bulgaria\|ep\|6510 | | human | | + | $\mathcal{N}(\mu = 6510, \sigma = 24)$ |
| KAP002\|Russian_Federation\|ep\|3513 | | human | | + | $\mathcal{U}(3213, 3813)$ |
| MIB011\|Czech_Republic\|ep\|3825 | | human | | + | $\mathcal{N}(\mu = 3826, \sigma = 34)$ |
| BOO006\|Russian_Federation\|ep\|3533 | | human | | + | $\mathcal{U}(3493, 3573)$ |
| BOO008\|Russian_Federation\|ep\|3533 | | human | | + | $\mathcal{U}(3493, 3573)$ |
| CHT001\|Slovakia\|ep\|2213 | | human | | + | $\mathcal{U}(2163, 2263)$ |
| I1321\|Georgia\|ep\|1935 | | human | | + | $\mathcal{N}(\mu = 1936, \sigma = 25)$ |
| KBD002\|Russian_Federation\|ep\|4097 | | human | | + | $\mathcal{N}(\mu = 4097, \sigma = 27)$ |
| SGR004\|Russian_Federation\|ep\|5013 | | human | | + | $\mathcal{U}(4513, 5513)$ |
| CAO009\|Cuba\|ep\|1393 | | human | | + | $\mathcal{N}(\mu = 1393, \sigma = 15)$ |
| KUE033\|Peru\|ep\|644 | | human | | + | $\mathcal{N}(\mu = 644, \sigma = 24)$ |
| MAP007\|Peru\|ep\|581 | | human | | + | $\mathcal{N}(\mu = 581, \sigma = 7)$ |
| PLS004\|Peru\|ep\|495 | | human | | + | $\mathcal{U}(453, 538)$ |
| SJN013\|Mexico\|ep\|585 | | human | | + | $\mathcal{N}(\mu = 586, \sigma = 6)$ |
| SGR003\|Russian_Federation\|en\|3913 | | human | | - | $\mathcal{U}(3813, 4013)$ |
| VLI060\|Czech_Republic\|ep\|3863 | | human | | + | $\mathcal{U}(3713, 4013)$ |

*Continued on next page*

| Sequence | Accession | Host | Genotype | HBeAg | Age |
|---|---|---|---|---|---|
| MK5004\|Russian_Federation\|en\|5229 | | human | | - | $\mathcal{N}(\mu = 5229, \sigma = 63)$ |
| MK5009\|Russian_Federation\|en\|4772 | | human | | - | $\mathcal{N}(\mu = 4772, \sigma = 59)$ |
| PDA003\|Czech_Republic\|en\|4183 | | human | | - | $\mathcal{N}(\mu = 4183, \sigma = 52)$ |
| LEU065\|Germany\|en\|4071 | | human | | - | $\mathcal{N}(\mu = 4072, \sigma = 41)$ |
| AF222323\|NA\|Chimp\|ep\|13 | AF222323 | Chimp | | + | 13 |
| AB032433\|LBR\|Chimp\|en\|31 | AB032433 | Chimp | | - | 31 |
| AY330911\|CMR\|Chimp\|ep\|10 | AY330911 | Chimp | | + | 10 |
| AJ131567\|CMR\|Gorilla\|ep\|14 | AJ131567 | Gorilla | | + | 14 |
| FM209516\|TWN\|Gibbon\|ep\|7 | FM209516 | Gibbon | | + | 7 |
| U46935\|NA\|Gibbon\|ep\|23 | U46935 | Gibbon | | + | 23 |
| AJ131571\|GER\|Gibbon\|ep\|14 | AJ131571 | Gibbon | | + | 14 |
| AY781180\|KHM\|Gibbon\|ep\|9 | AY781180 | Gibbon | | + | 9 |
| EU155824\|THA\|Orang\|ep\|6 | EU155824 | Orang | | + | 6 |
| AF193863\|NA\|Orang\|ep\|14 | AF193863 | Orang | | + | 14 |
| 11KBM13\|China\|ep\|3468 | | human | | + | $\mathcal{N}(\mu = 3468, \sigma = 42)$ |
| 91KLH18\|China\|ep\|2845 | | human | | + | $\mathcal{N}(\mu = 2845, \sigma = 41)$ |
| 98JJLM9\|China\|ep\|4128 | | human | | + | $\mathcal{N}(\mu = 4128, \sigma = 40)$ |
| ZQM16\|China\|ep\|1643 | | human | | + | $\mathcal{N}(\mu = 1643, \sigma = 34)$ |
| XHM18\|China\|ep\|2503 | | human | | + | $\mathcal{N}(\mu = 2503, \sigma = 36)$ |
| XBQM47\|China\|ep\|2958 | | human | | + | $\mathcal{N}(\mu = 2958, \sigma = 37)$ |
| XBQM86\|China\|ep\|2958 | | human | | + | $\mathcal{N}(\mu = 2958, \sigma = 37)$ |
| XBQM125\|China\|ep\|2958 | | human | | + | $\mathcal{N}(\mu = 2958, \sigma = 37)$ |
| FLTM101\|China\|ep\|2218 | | human | | + | $\mathcal{N}(\mu = 2218, \sigma = 31)$ |
| FLTM48\|China\|ep\|2193 | | human | | + | $\mathcal{N}(\mu = 2193, \sigma = 35)$ |
| MY12\|Mongolia\|ep\|778 | | human | | + | $\mathcal{N}(\mu = 778, \sigma = 28)$ |
| MY17\|Mongolia\|ep\|778 | | human | | + | $\mathcal{N}(\mu = 778, \sigma = 28)$ |
| AT19\|Mongolia\|ep\|2283 | | human | | + | $\mathcal{N}(\mu = 2283, \sigma = 39)$ |
| AT24\|Mongolia\|ep\|2283 | | human | | + | $\mathcal{N}(\mu = 2283, \sigma = 39)$ |
| AT7\|Mongolia\|ep\|2283 | | human | | + | $\mathcal{N}(\mu = 2283, \sigma = 39)$ |
| TJZM25_2\|China\|ep\|1828 | | human | | + | $\mathcal{N}(\mu = 1828, \sigma = 38)$ |
| XBQM20\|China\|ep\|2948 | | human | | + | $\mathcal{N}(\mu = 2948, \sigma = 37)$ |
| XBQM46\|China\|ep\|2948 | | human | | + | $\mathcal{N}(\mu = 2948, \sigma = 37)$ |
| XN12\|Russia\|ep\|2123 | | human | | + | $\mathcal{N}(\mu = 2123, \sigma = 31)$ |
| FLTM97\|China\|ep\|2193 | | human | | + | $\mathcal{N}(\mu = 2193, \sigma = 35)$ |
| XHM12\|China\|ep\|2503 | | human | | + | $\mathcal{N}(\mu = 2503, \sigma = 36)$ |
| HHM29\|China\|ep\|463 | | human | | + | $\mathcal{N}(\mu = 463, \sigma = 29)$ |
| JHM2098\|China\|ep\|2563 | | human | | + | $\mathcal{N}(\mu = 2563, \sigma = 37)$ |
| 96NVZIM6\|China\|ep\|5063 | | human | | + | $\mathcal{N}(\mu = 5063, \sigma = 47)$ |
| SBSM101\|China\|ep\|1563 | | human | | + | $\mathcal{N}(\mu = 1563, \sigma = 34)$ |

Table 2: HBV recombination events. For each event, we list breakpoint positions in the multiple sequence alignment, recombinant sequences, and inferred parental sequences. BP = breakpoint. # = The actual breakpoint position is undetermined (e.g., obscured by subsequent recombination or outside the analysed sequence region). * = The recombinant sequence may have been misidentified (i.e., one of the inferred parents could be recombinant). Minor parent = parent contributing the smaller fraction of the sequence. Major parent = parent contributing the larger fraction of the sequence. Unknown = a missing parental sequence inferred from the alignment. NS = no statistically significant P-value was obtained for the recombination event.

<table>
<tr><th>Event</th><th>BP start</th><th>BP end</th><th>Recombinant sequences</th><th>Minor parent</th><th>Major parent</th></tr>
<tr><td>1</td><td>1759</td><td>2210</td><td>FJ386636|China|B2|ep|5</td><td>AY247031|Korea|C|ep|16</td><td>AB287326|Japan|B1|en|7</td></tr>
<tr><td></td><td></td><td></td><td>FJ386584|China|B2|ep|5</td><td></td><td></td></tr>
<tr><td></td><td></td><td></td><td>FJ386600|China|B2|ep|5</td><td></td><td></td></tr>
<tr><td></td><td></td><td></td><td>GQ924653|Malaysia|B2|ep|6</td><td></td><td></td></tr>
<tr><td>2</td><td>1563</td><td>2037</td><td>*SGR003|Russian_Federation|en|3913</td><td>UZZ099|Italy_Sicily|en|6011</td><td>KAP002|Russian_Federation|ep|3513</td></tr>
<tr><td>3</td><td>1483</td><td>2212</td><td>AB194947|Cameroon|E|ep|19</td><td>KBD002|Russian_Federation|ep|4097</td><td>ELT006|Spain|en|5863</td></tr>
<tr><td></td><td></td><td></td><td>AB205129|Ghana|E|ep|13</td><td>RISE386|Russian_Federation|ep|4184</td><td>Karsdorf|Germany|en|7089</td></tr>
<tr><td></td><td></td><td></td><td>AB205188|Ghana|E|epi|13</td><td>BOO006|Russian_Federation|ep|3533</td><td>HAL05|Germany|en|7105</td></tr>
<tr><td></td><td></td><td></td><td>AB205189|Ghana|E|ep|13</td><td>BOO008|Russian_Federation|ep|3533</td><td>UZZ099|Italy_Sicily|en|6011</td></tr>
<tr><td></td><td></td><td></td><td>AB205190|Ghana|E|ep|13</td><td>SGR004|Russian_Federation|ep|5013</td><td>MK5004|Russian_Federation|en|5229</td></tr>
<tr><td></td><td></td><td></td><td>AB205191|Ghana|E|ep|13</td><td></td><td></td></tr>
<tr><td></td><td></td><td></td><td>AB205192|Ghana|E|ep|13</td><td></td><td></td></tr>
<tr><td></td><td></td><td></td><td>AP007262|Zambia|E|ep|12</td><td></td><td></td></tr>
<tr><td></td><td></td><td></td><td>DQ060822|Angola|E|ep|20</td><td></td><td></td></tr>
<tr><td></td><td></td><td></td><td>DQ060823|Angola|E|ep|20</td><td></td><td></td></tr>
<tr><td>4</td><td>1615#</td><td>1728</td><td>ISB002|Italy_Sardinia|en|3688</td><td>DER027|Germany|ep|4013</td><td>ELT006|Spain|en|5863</td></tr>
<tr><td>5</td><td>1303</td><td>1462#</td><td>*HQ700521|PNG|C|ep|9</td><td>AF223964|Argentina|F|ep|17</td><td>BOO008|Russian_Federation|ep|3533</td></tr>
<tr><td></td><td></td><td></td><td>HQ700519|PNG|C|ep|10</td><td>AF223963|Argentina|F|ep|17</td><td></td></tr>
<tr><td></td><td></td><td></td><td>HQ700522|PNG|C|ep|9[T]</td><td>AY179735|Argintina|F|ep|16</td><td></td></tr>
<tr><td></td><td></td><td></td><td></td><td>AB064316|USA|F|en|17</td><td></td></tr>
<tr><td>6</td><td>2321</td><td>2492</td><td>*FM209516|TWN|Gibbon|ep|7</td><td>AF223965|Argentina|F|ep|17</td><td>Unknown(AB059660|USA|H|ep|12)</td></tr>
<tr><td></td><td></td><td></td><td></td><td>AF223962|Argentina|F|ep|17</td><td></td></tr>
<tr><td>7</td><td>1445#</td><td>2574</td><td>ELT006|Spain|en|5863</td><td>MK5009|Russian_Federation|en|4772</td><td>Unknown(CLL005|Spain|ep|4913)</td></tr>
<tr><td></td><td></td><td></td><td>I0784|Bulgaria|en|6313[P]</td><td></td><td></td></tr>
<tr><td>8</td><td>3129</td><td>1444</td><td>*SID005|Italy_Sardinia|en|6157</td><td>ELT005|Spain|ep|7227</td><td>Unknown(ARM001|Armenia|ep|5511)</td></tr>
<tr><td>9</td><td>1568</td><td>1742</td><td>*AB064312|USA|G|en|11</td><td>Unknown(AB113878|Japan|C|en|13)</td><td>RISE387|Russian_Federation|en|4278</td></tr>
<tr><td></td><td></td><td></td><td>AB056513|USA|G|en|11</td><td>Unknown(AF223954|EastAsia|C|ep|25)</td><td></td></tr>
<tr><td></td><td></td><td></td><td>AF405706|Germany|G|en|11</td><td>Unknown(AF223955|EastAsia|C|ep|24)</td><td></td></tr>
<tr><td></td><td></td><td></td><td></td><td>Unknown(AF223957|EastAsia|C|ep|27)</td><td></td></tr>
<tr><td></td><td></td><td></td><td></td><td>Unknown(AF223956|EastAsia|C|ep|26)</td><td></td></tr>
<tr><td></td><td></td><td></td><td></td><td>Unknown(AF223960|EastAsia|C|ep|28)</td><td></td></tr>
<tr><td></td><td></td><td></td><td></td><td>Unknown(AF223961|EastAsia|C|ep|22)</td><td></td></tr>
<tr><td></td><td></td><td></td><td></td><td>Unknown(AY247031|Korea|C|ep|16)</td><td></td></tr>
<tr><td></td><td></td><td></td><td></td><td>Unknown(AY167092|Taiwan|C|epi|16)</td><td></td></tr>
<tr><td></td><td></td><td></td><td></td><td>Unknown(HQ700505|VANUATA|C|ep|9)</td><td></td></tr>
<tr><td></td><td></td><td></td><td></td><td>Unknown(HQ700506|N_CALEDONIA|C|ep|10)</td><td></td></tr>
<tr><td></td><td></td><td></td><td></td><td>Unknown(HQ700507|N_CALEDONIA|C|ep|12)</td><td></td></tr>
<tr><td></td><td></td><td></td><td></td><td>Unknown(HQ700508|N_CALEDONIA|C|ep|10)</td><td></td></tr>
<tr><td></td><td></td><td></td><td></td><td>Unknown(HQ700509|N_CALEDONIA|C|ep|10)</td><td></td></tr>
<tr><td></td><td></td><td></td><td></td><td>Unknown(HQ700527|PNG|C|ep|9)</td><td></td></tr>
</table>

*Continued on next page*

<table>
<tr><th>Event</th><th>BP start</th><th>BP end</th><th>Recombinant sequences</th><th>Minor parent</th><th>Major parent</th></tr>
<tr><td></td><td></td><td></td><td></td><td>Unknown(HQ700519|PNG|C|ep|10)</td><td></td></tr>
<tr><td></td><td></td><td></td><td></td><td>Unknown(HQ700520|PNG|C|ep|9)</td><td></td></tr>
<tr><td></td><td></td><td></td><td></td><td>Unknown(HQ700522|PNG|C|ep|9)</td><td></td></tr>
<tr><td></td><td></td><td></td><td></td><td>Unknown(HQ700530|PNG|C|ep|19)</td><td></td></tr>
<tr><td></td><td></td><td></td><td></td><td>Unknown(HQ700485|NZL|C|ep|29)</td><td></td></tr>
<tr><td></td><td></td><td></td><td></td><td>Unknown(HQ700461|NZL|C|ep|29)</td><td></td></tr>
<tr><td></td><td></td><td></td><td></td><td>Unknown(HQ700526|PNG|C|ep|8)</td><td></td></tr>
<tr><td></td><td></td><td></td><td></td><td>Unknown(HQ700462|NZL|C|en|17)</td><td></td></tr>
<tr><td></td><td></td><td></td><td></td><td>Unknown(HQ700490|NZL|C|en|20)</td><td></td></tr>
<tr><td>10</td><td>2875</td><td>692</td><td>*MK5009|Russian_Federation|en|4772</td><td>PDA001|Czech_Republic|ep|4069</td><td>Unknown (DQ020003|UnitedArabEmirates|A|ep|25)</td></tr>
<tr><td></td><td></td><td></td><td>MK5004|Russian_Federation|en|5229</td><td></td><td>Unknown(AB194951|Cameroon|A|ep|19)</td></tr>
<tr><td></td><td></td><td></td><td></td><td></td><td>Unknown(AB194952|Cameroon|A|ep|19)</td></tr>
<tr><td></td><td></td><td></td><td></td><td></td><td>Unknown(AY934766|Somalia|A|ep|18)</td></tr>
<tr><td></td><td></td><td></td><td></td><td></td><td>Unknown(AY934767|Somalia|A|ep|17)</td></tr>
<tr><td></td><td></td><td></td><td></td><td></td><td>Unknown(AY934768|Somalia|A|ep|18)</td></tr>
<tr><td></td><td></td><td></td><td></td><td></td><td>Unknown(AY934769|Somalia|A|ep|18)</td></tr>
<tr><td></td><td></td><td></td><td></td><td></td><td>Unknown(AY934770|Somalia|A|ep|17)</td></tr>
<tr><td></td><td></td><td></td><td></td><td></td><td>Unknown(AY934771|Somalia|A|ep|17)</td></tr>
<tr><td></td><td></td><td></td><td></td><td></td><td>Unknown(AY934772|Uganda|A|ep|12)</td></tr>
<tr><td></td><td></td><td></td><td></td><td></td><td>Unknown(AY934774|Philippines|A|ep|17)</td></tr>
<tr><td></td><td></td><td></td><td></td><td></td><td>Unknown(DQ020002|Congo|A|ep|10)</td></tr>
<tr><td></td><td></td><td></td><td></td><td></td><td>Unknown(FJ692556|Nigeria|A3|ep|9)</td></tr>
<tr><td></td><td></td><td></td><td></td><td></td><td>Unknown(FJ692593|Haiti|A3|ep|7)</td></tr>
<tr><td></td><td></td><td></td><td></td><td></td><td>Unknown(FJ692594|Haiti|A3|ep|7)</td></tr>
<tr><td></td><td></td><td></td><td></td><td></td><td>Unknown(FJ692595|Haiti|A3|ep|7)</td></tr>
<tr><td></td><td></td><td></td><td></td><td></td><td>Unknown(FJ692596|Haiti|A3|ep|7)</td></tr>
<tr><td></td><td></td><td></td><td></td><td></td><td>Unknown(FJ692597|Haiti|A3|ep|7)</td></tr>
<tr><td></td><td></td><td></td><td></td><td></td><td>Unknown(FJ692598|Haiti|A3|ep|7)</td></tr>
<tr><td></td><td></td><td></td><td></td><td></td><td>Unknown(FJ692599|Haiti|A3|ep|7)</td></tr>
<tr><td></td><td></td><td></td><td></td><td></td><td>Unknown(FJ692600|Haiti|A3|ep|7)</td></tr>
<tr><td></td><td></td><td></td><td></td><td></td><td>Unknown(FJ692602|Haiti|A3|ep|7)</td></tr>
<tr><td></td><td></td><td></td><td></td><td></td><td>Unknown(FJ692603|Haiti|A3|ep|7)</td></tr>
<tr><td></td><td></td><td></td><td></td><td></td><td>Unknown(FJ692604|Haiti|A3|ep|7)</td></tr>
<tr><td></td><td></td><td></td><td></td><td></td><td>Unknown(FJ692605|Haiti|A3|ep|7)</td></tr>
<tr><td></td><td></td><td></td><td></td><td></td><td>Unknown(FJ692606|Haiti|A3|ep|7)</td></tr>
<tr><td></td><td></td><td></td><td></td><td></td><td>Unknown(FJ692607|Haiti|A3|ep|7)</td></tr>
<tr><td></td><td></td><td></td><td></td><td></td><td>Unknown(FJ692608|Haiti|A3|ep|7)</td></tr>
<tr><td></td><td></td><td></td><td></td><td></td><td>Unknown(FJ692609|Haiti|A3|en|7)</td></tr>
<tr><td></td><td></td><td></td><td></td><td></td><td>Unknown(FJ692611|Haiti|A3|ep|7)</td></tr>
<tr><td></td><td></td><td></td><td></td><td></td><td>Unknown(FJ692613|Haiti|A3|ep|7)</td></tr>
<tr><td></td><td></td><td></td><td></td><td></td><td>Unknown(DA119|Slovakia|ep|1563)</td></tr>
<tr><td></td><td></td><td></td><td></td><td></td><td>Unknown(SJN001|Mexico|A3|ep|473)</td></tr>
<tr><td></td><td></td><td></td><td></td><td></td><td>Unknown(CHT001|Slovakia|ep|2213)</td></tr>
<tr><td></td><td></td><td></td><td></td><td></td><td>Unknown(I1321|Georgia|ep|1935)</td></tr>
<tr><td>11</td><td>1881</td><td>2244</td><td>91KLH18|China|ep|2845</td><td>Unknown (FJ692605|Haiti|A3|ep|7)</td><td>KP659219|Alaska|B5|en|7</td></tr>
<tr><td></td><td></td><td></td><td>XHM18|China|ep|2503</td><td>Unknown(AB287318|Alaska|B5|en|9)</td><td>FJ692605|Haiti|A3|ep|7</td></tr>
<tr><td></td><td></td><td></td><td>XBQM47|China|ep|2958</td><td>Unknown(KP659219|Alaska|B5|en|7)</td><td></td></tr>
<tr><td></td><td></td><td></td><td>FLTM101|China|ep|2218[T]</td><td></td><td></td></tr>
</table>

*Continued on next page*

<table>
<tr><th>Event</th><th>BP start</th><th>BP end</th><th>Recombinant sequences</th><th>Minor parent</th><th>Major parent</th></tr>
<tr><td></td><td></td><td></td><td>FLTM48|China|ep|2193[T]</td><td></td><td></td></tr>
<tr><td></td><td></td><td></td><td>AT24|Mongolia|ep|2283[T]</td><td></td><td></td></tr>
<tr><td></td><td></td><td></td><td>AT7|Mongolia|ep|2283[T]</td><td></td><td></td></tr>
<tr><td></td><td></td><td></td><td>XN12|Russia|ep|2123[T]</td><td></td><td></td></tr>
<tr><td></td><td></td><td></td><td>FLTM97|China|ep|2193[T]</td><td></td><td></td></tr>
<tr><td></td><td></td><td></td><td>XHM12|China|ep|2503[T]</td><td></td><td></td></tr>
<tr><td></td><td></td><td></td><td>JHM2098|China|ep|2563[T]</td><td></td><td></td></tr>
<tr><td></td><td></td><td></td><td>96NVZIM6|China|ep|5063[T]</td><td></td><td></td></tr>
<tr><td>12</td><td>1609</td><td>2186</td><td>*KP659221|Alaska|B5|en|8</td><td>Unknown (AB287324|WestGreenland|B5|en|9)</td><td>AB287317|Alaska|B5|en|39</td></tr>
<tr><td></td><td></td><td></td><td></td><td>Unknown(AB287317|Alaska|B5|en|39)</td><td>AB287324|WestGreenland|B5|en|9</td></tr>
<tr><td>13</td><td>1486</td><td>2212</td><td>*KBD002|Russian_Federation|ep|4097</td><td>DA51|Kyrgyzstan|ep|2293</td><td>Unknown (UZZ099|Italy_Sicily|en|6011)</td></tr>
<tr><td></td><td></td><td></td><td>RISE386|Russian_Federation|ep|4184</td><td>AF043593|Germany|D|ep|24</td><td>Unknown(VLI060|Czech_Republic|ep|3863)</td></tr>
<tr><td></td><td></td><td></td><td>BOO006|Russian_Federation|ep|3533[P]</td><td>AY741797|Iran|D|ep|10</td><td>Unknown(MK5004|Russian_Federation|en|5229)</td></tr>
<tr><td></td><td></td><td></td><td>BOO008|Russian_Federation|ep|3533</td><td>AB090268|Japan|D|ep|16</td><td></td></tr>
<tr><td></td><td></td><td></td><td>SGR004|Russian_Federation|ep|5013[P]</td><td>AB090269|Japan|D|ep|21</td><td></td></tr>
<tr><td></td><td></td><td></td><td>XBQM86|China|ep|2958</td><td>AB078033|Japan|D|ep|15</td><td></td></tr>
<tr><td></td><td></td><td></td><td></td><td>AB090270|Japan|D|ep|14</td><td></td></tr>
<tr><td></td><td></td><td></td><td></td><td>AB109475|Japan|D|ep|12</td><td></td></tr>
<tr><td></td><td></td><td></td><td></td><td>AB109476|Japan|D|ep|16</td><td></td></tr>
<tr><td></td><td></td><td></td><td></td><td>AB120308|Japan|D|ep|31</td><td></td></tr>
<tr><td></td><td></td><td></td><td></td><td>AF121240|NA|D|ep|19</td><td></td></tr>
<tr><td></td><td></td><td></td><td></td><td>HQ700497|FIJI|D|ep|9</td><td></td></tr>
<tr><td></td><td></td><td></td><td></td><td>HQ700503|TONGA|D|ep|21</td><td></td></tr>
<tr><td></td><td></td><td></td><td></td><td>HQ700501|SAMOA|D|ep|22</td><td></td></tr>
<tr><td></td><td></td><td></td><td></td><td>HQ700511|N_CALEDONIA|D|ep|12</td><td></td></tr>
<tr><td></td><td></td><td></td><td></td><td>HQ700512|N_CALEDONIA|D|ep|12</td><td></td></tr>
<tr><td></td><td></td><td></td><td></td><td>HQ700513|N_CALEDONIA|D|ep|12</td><td></td></tr>
<tr><td></td><td></td><td></td><td></td><td>HQ700514|N_CALEDONIA|D|ep|12</td><td></td></tr>
<tr><td></td><td></td><td></td><td></td><td>HQ700524|PNG|D|ep|9</td><td></td></tr>
<tr><td></td><td></td><td></td><td></td><td>HQ700525|PNG|D|ep|9</td><td></td></tr>
<tr><td></td><td></td><td></td><td></td><td>HQ700541|Kiribati|D|ep|8</td><td></td></tr>
<tr><td></td><td></td><td></td><td></td><td>HQ700538|Kiribati|D|ep|9</td><td></td></tr>
<tr><td></td><td></td><td></td><td></td><td>HQ700464|NZL|D|ep|29</td><td></td></tr>
<tr><td></td><td></td><td></td><td></td><td>HQ700442|NZL|D|ep|14</td><td></td></tr>
<tr><td></td><td></td><td></td><td></td><td>HQ700466|NZL|D|ep|29</td><td></td></tr>
<tr><td></td><td></td><td></td><td></td><td>HQ700470|NZL|D|ep|14</td><td></td></tr>
<tr><td></td><td></td><td></td><td></td><td>HQ700474|NZL|D|ep|14</td><td></td></tr>
<tr><td></td><td></td><td></td><td></td><td>HQ700484|NZL|D|ep|29</td><td></td></tr>
<tr><td></td><td></td><td></td><td></td><td>HQ700459|NZL|D|ep|20</td><td></td></tr>
<tr><td></td><td></td><td></td><td></td><td>HQ700478|NZL|D|en|19</td><td></td></tr>
<tr><td></td><td></td><td></td><td></td><td>HQ700481|NZL|D|en|14</td><td></td></tr>
<tr><td></td><td></td><td></td><td></td><td>HQ700472|NZL|D|en|16</td><td></td></tr>
<tr><td></td><td></td><td></td><td></td><td>HQ700458|NZL|D|en|16</td><td></td></tr>
<tr><td></td><td></td><td></td><td></td><td>HQ700446|NZL|D|ep|14</td><td></td></tr>
<tr><td></td><td></td><td></td><td></td><td>HQ700448|NZL|D|ep|14</td><td></td></tr>
<tr><td></td><td></td><td></td><td></td><td>HQ700455|NZL|D|ep|14</td><td></td></tr>
<tr><td></td><td></td><td></td><td></td><td>AY741798|Iran|D|en|9</td><td></td></tr>
<tr><td></td><td></td><td></td><td></td><td>AB109478|Japan|D|en|15</td><td></td></tr>
</table>

*Continued on next page*

<table>
<tr><th>Event</th><th>BP start</th><th>BP end</th><th>Recombinant sequences</th><th>Minor parent</th><th>Major parent</th></tr>
<tr><td></td><td></td><td></td><td></td><td>AB109479|Japan|D|en|16</td><td></td></tr>
<tr><td></td><td></td><td></td><td></td><td>AB119251|Japan|D|en|11</td><td></td></tr>
<tr><td></td><td></td><td></td><td></td><td>AB119252|Japan|D|en|11</td><td></td></tr>
<tr><td></td><td></td><td></td><td></td><td>AB119254|Japan|D|en|10</td><td></td></tr>
<tr><td></td><td></td><td></td><td></td><td>AF121239|Japan|D|en|20</td><td></td></tr>
<tr><td></td><td></td><td></td><td></td><td>HQ700540|Kiribati|D|en|9</td><td></td></tr>
<tr><td></td><td></td><td></td><td></td><td>HQ700533|Kiribati|D|en|9</td><td></td></tr>
<tr><td></td><td></td><td></td><td></td><td>HQ700534|Kiribati|D|en|9</td><td></td></tr>
<tr><td></td><td></td><td></td><td></td><td>HQ700535|Kiribati|D|en|9</td><td></td></tr>
<tr><td></td><td></td><td></td><td></td><td>HQ700537|Kiribati|D|en|9</td><td></td></tr>
<tr><td></td><td></td><td></td><td></td><td>HQ700536|Kiribati|D|en|9</td><td></td></tr>
<tr><td></td><td></td><td></td><td></td><td>HQ700510|N_CALEDONIA|D|en|10</td><td></td></tr>
<tr><td></td><td></td><td></td><td></td><td>HQ700492|Fiji|D|en|9</td><td></td></tr>
<tr><td></td><td></td><td></td><td></td><td>HQ700493|Fiji|D|en|9</td><td></td></tr>
<tr><td></td><td></td><td></td><td></td><td>HQ700494|Fiji|D|en|9</td><td></td></tr>
<tr><td></td><td></td><td></td><td></td><td>HQ700500|Samoa|D|en|20</td><td></td></tr>
<tr><td></td><td></td><td></td><td></td><td>DA29|Kazakhstan|ep|818</td><td></td></tr>
<tr><td></td><td></td><td></td><td></td><td>DA222|Kazakhstan|ep|1163</td><td></td></tr>
<tr><td></td><td></td><td></td><td></td><td>MG585269|Italy|D3|ep|444</td><td></td></tr>
<tr><td></td><td></td><td></td><td></td><td>Petersberg|Germany|ep|931</td><td></td></tr>
<tr><td></td><td></td><td></td><td></td><td>AKB003|Kazakhstan|ep|2649</td><td></td></tr>
<tr><td></td><td></td><td></td><td></td><td>BRE008|Kazakhstan|ep|1701</td><td></td></tr>
<tr><td></td><td></td><td></td><td></td><td>BRE026|Kazakhstan|ep|2363</td><td></td></tr>
<tr><td></td><td></td><td></td><td></td><td>BRE028|Kazakhstan|ep|2363</td><td></td></tr>
<tr><td></td><td></td><td></td><td></td><td>I0157|Great_Britain|ep|1295</td><td></td></tr>
<tr><td></td><td></td><td></td><td></td><td>I0161|Great_Britain|ep|1228</td><td></td></tr>
<tr><td></td><td></td><td></td><td></td><td>I0216|Italy|ep|2663</td><td></td></tr>
<tr><td></td><td></td><td></td><td></td><td>I1344|Russian_Federation|ep|713</td><td></td></tr>
<tr><td></td><td></td><td></td><td></td><td>KIL044|Ireland|en|838</td><td></td></tr>
<tr><td></td><td></td><td></td><td></td><td>KRA001|Germany|ep|797</td><td></td></tr>
<tr><td></td><td></td><td></td><td></td><td>KRA010|Germany|ep|683</td><td></td></tr>
<tr><td></td><td></td><td></td><td></td><td>MAY017|Russian_Federation|ep|663</td><td></td></tr>
<tr><td></td><td></td><td></td><td></td><td>MLR005|Russian_Federation|ep|713</td><td></td></tr>
<tr><td></td><td></td><td></td><td></td><td>OAI017|Great_Britain|ep|1513</td><td></td></tr>
<tr><td></td><td></td><td></td><td></td><td>ORE002|Russian_Federation|ep|2585</td><td></td></tr>
<tr><td></td><td></td><td></td><td></td><td>RMI002|Germany|ep|613</td><td></td></tr>
<tr><td></td><td></td><td></td><td></td><td>SHK001|Russian_Federation|ep|811</td><td></td></tr>
<tr><td></td><td></td><td></td><td></td><td>STR144|Belgium|ep|469</td><td></td></tr>
<tr><td></td><td></td><td></td><td></td><td>ZWE008|Switzerland|ep|680</td><td></td></tr>
<tr><td></td><td></td><td></td><td></td><td>XBQM46|China|ep|2948</td><td></td></tr>
<tr><td>14</td><td>2689</td><td>878</td><td>*RISE563|Germany|ep|4484</td><td>RISE254|Hungary|ep|4005</td><td>Unknown(HOP004|Czech_Republic|ep|4310)</td></tr>
<tr><td>15</td><td>2092</td><td>2849</td><td>SGR003|Russian_Federation|en|3913</td><td>PDA001|Czech_Republic|ep|4069</td><td>KAP002|Russian_Federation|ep|3513</td></tr>
<tr><td>16</td><td>1566#</td><td>2207</td><td>*HAL05|Germany|en|7105<br>Karsdorf|Germany|en|7089</td><td>Unknown(RISE387|Russian_Federation|en|4278)</td><td>I0411|Spain|ep|7192</td></tr>
<tr><td>17</td><td>2034</td><td>2207</td><td>RISE387|Russian_Federation|en|4278</td><td>DA51|Kyrgyzstan|ep|2293</td><td>HAL05|Germany|en|7105</td></tr>
</table>
